# A Chloroplast Protein Atlas Reveals Novel Structures and Spatial Organization of Biosynthetic Pathways

**DOI:** 10.1101/2022.05.31.493820

**Authors:** Lianyong Wang, Weronika Patena, Kelly A. Van Baalen, Yihua Xie, Emily R. Singer, Sophia Gavrilenko, Michelle Warren-Williams, Linqu Han, Henry R. Harrigan, Vivian Chen, Vinh T.N.P. Ton, Saw Kyin, Henry H. Shwe, Matthew H. Cahn, Alexandra T. Wilson, Jianping Hu, Danny J. Schnell, Claire D. McWhite, Martin Jonikas

## Abstract

Chloroplasts are eukaryotic photosynthetic organelles that drive the global carbon cycle. Despite their importance, our understanding of their protein composition, function, and spatial organization remains limited. Here, we determined the localizations of 1,032 candidate chloroplast proteins by using fluorescent protein tagging in the model alga *Chlamydomonas reinhardtii*. The localizations provide insights into the functions of hundreds of poorly-characterized proteins, including identifying novel components of nucleoids, plastoglobules, and the pyrenoid. We discovered and further characterized novel organizational features, including eleven chloroplast punctate structures, cytosolic crescent structures, and diverse unexpected spatial distributions of enzymes within the chloroplast. We observed widespread protein targeting to multiple organelles, identifying proteins that likely function in multiple compartments. We also used machine learning to predict the localizations of all *Chlamydomonas* proteins. The strains and localization atlas developed here will serve as a resource to enable studies of chloroplast architecture and functions.

**Graphical Abstract:** 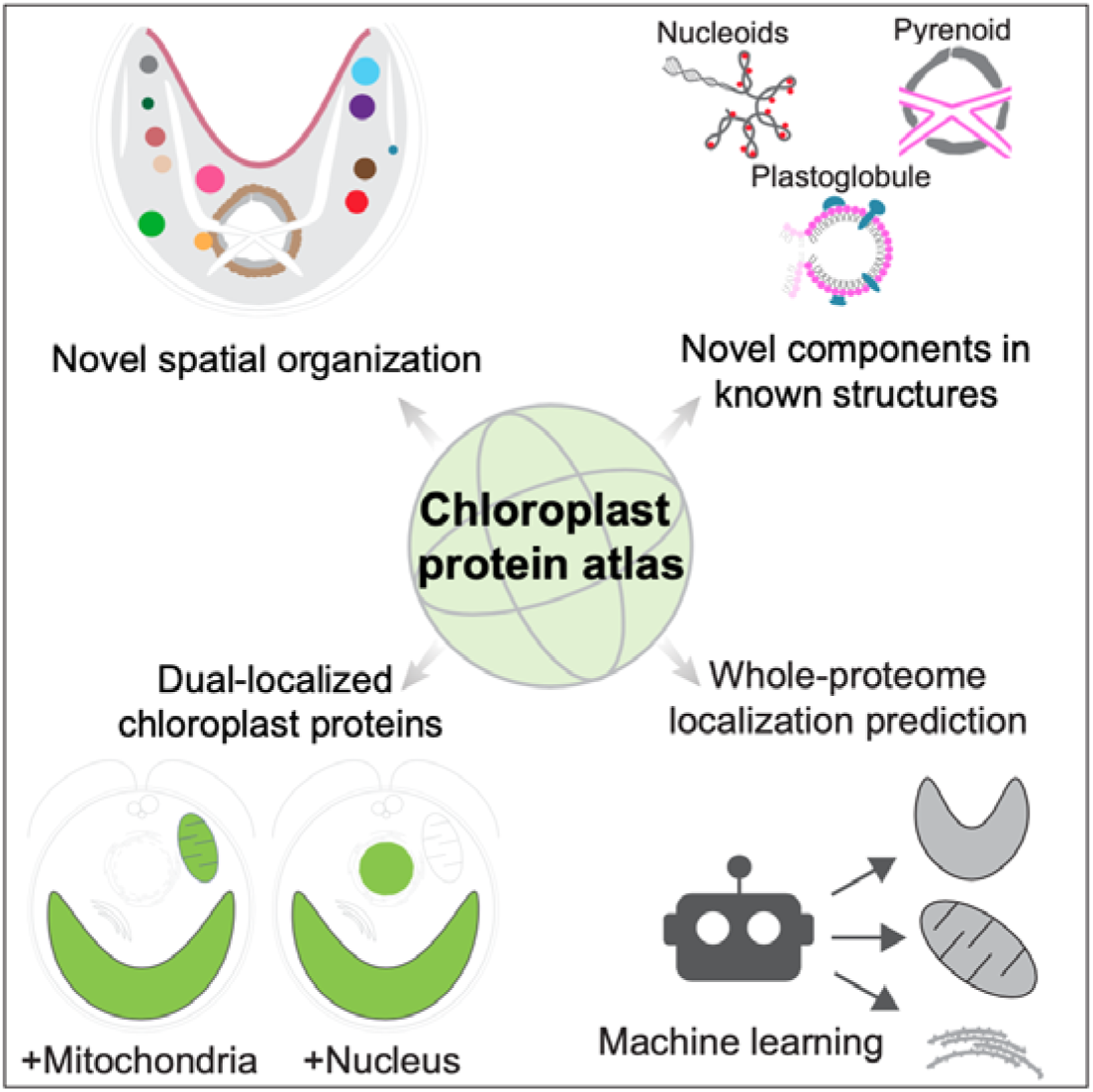

**Highlights:**

- 1,032 candidate chloroplast proteins localized by fluorescent tagging.
- This protein atlas reveals novel chloroplast structures, functional regions, and components.
- Prevalent dual-organelle localization suggests extensive cross-compartment coordination.
- Atlas-trained machine learning predicts localizations of all *C. reinhardtii* proteins.

## INTRODUCTION

The chloroplast is a hallmark organelle of eukaryotic photosynthetic organisms. Over 85% of global biological light energy capture, CO_2_ fixation, and O_2_ production happens in a chloroplast, driving the biochemistry of the biosphere (Behrenfeld et al., 2001; Rousseaux and Gregg, 2013). In addition to performing photosynthesis, the chloroplast plays essential roles in key cellular processes including amino acid synthesis (Hildebrandt et al., 2015), starch synthesis (Pfister and Zeeman, 2016), lipid metabolism (Hölzl and Dörmann, 2019), isoprenoid synthesis (Lange et al., 2000), purine/pyrimidine synthesis (Zrenner et al., 2006), and the immune response of land plants (Nomura et al., 2012). Despite the chloroplast’s importance, the mechanisms of its function and regulation are still not well understood.

All chloroplasts are thought to have originated from a single primary endosymbiosis of a free-living photosynthetic cyanobacterium by a host eukaryotic cell (Keeling, 2010). Over time, the engulfed cyanobacterium lost its autonomy and transferred most of its genes to the nucleus. In parallel, sophisticated communication evolved between the chloroplast and other organelles. This endosymbiosis event is thought to have given rise to the *Archaeplastidia* eukaryotic supergroup, which includes land plants, red algae, and green algae. Secondary endosymbiosis of members of *Archaeplastidia* then produced the more complex chloroplasts found in other eukaryotic supergroups including the coccolithophores and diatoms. Hereafter, we focus on the chloroplast of *Archaeplastidia*, which remain dominant on a global scale, with land plants performing the vast majority of photosynthesis on land and green algae contributing a significant portion of photosynthesis in the oceans (Behrenfeld et al., 2001; Rousseaux and Gregg, 2013).

To understand the function and regulation of the chloroplast, we need to study its proteins and its sub-organellar organization. While the chloroplast has retained a minimal genome, the vast majority of its proteins are now encoded in the nucleus and are imported into the chloroplast (Leister, 2003). Although hundreds of nuclear-encoded proteins have recently been associated with the chloroplast through proteomics (Ferro et al., 2010; Terashima et al., 2010), phylogenetics (Karpowicz et al., 2011), and bioinformatics studies (Emanuelsson et al., 1999; Tardif et al., 2012), the protein composition of the chloroplast remains poorly defined. Moreover, most chloroplast-associated proteins remain functionally uncharacterized (Hooper et al., 2017; Leister, 2003; Leister and Kleine, 2008).

One promising starting point for understanding the functions of chloroplast-associated proteins is the systematic determination of their cellular and sub-chloroplast localizations (Huh et al., 2003; Thul et al., 2017). Existing knowledge of the chloroplast indicates that its functions are highly spatially organized into distinct regions within the organelle (Engel et al., 2015). For example, sub-chloroplast regions called nucleoids contain the chloroplast’s DNA (Kobayashi et al., 2002), chloroplast-traversing thylakoid membranes specialize in the photosynthetic capture of light energy (Wise and Hoober, 2006), and thylakoid membrane-associated lipid droplets called plastoglobules play roles in lipid metabolism (Ytterberg et al., 2006). Localizing a protein of unknown function to such a functionally-specialized region would immediately suggest a corresponding function for the protein.

Of the various approaches for determining subcellular protein localization, fluorescent protein tagging provides significant advantages for the present application. Fluorescent protein tagging is more accurate and has higher spatial resolution than proteome analysis of purified organelles or bioinformatics-based localization predictions, and can also reveal localizations and sub-organellar organization that were not previously known to exist (Mackinder et al., 2017). Furthermore, tagged strains can be affinity-purified and subjected to mass spectrometry-based proteomics to identify associating proteins that are other components of novel cellular structures.

To date, only a small subset of chloroplast proteins have been localized using fluorescent tagging or immunofluorescence. A recent comprehensive survey (Hooper et al., 2017) found that altogether only 582 of the ∼3,000 bioinformatically-predicted chloroplast proteins (Emanuelsson et al., 1999; Tardif et al., 2012) (∼19%) have been experimentally localized in the leading model land plant Arabidopsis. These numbers suggest that many opportunities lie ahead for the discovery of novel chloroplast structures and protein functions through systematic localization of fluorescently-tagged proteins.

The green alga *Chlamydomonas reinhardtii* (Chlamydomonas hereafter, Figure 1A) is a powerful model system for studying the cell biology of photosynthetic eukaryotes. Its unicellular nature and microbial lifestyle allow for higher throughput than land plant model systems, enabling systematic large-scale analysis of gene and protein function (Fauser et al., 2022). As an evolutionary relative of land plants (Gutman and Niyogi, 2004; Merchant et al., 2007), Chlamydomonas has been a critical model system that has revealed conserved pathways and key principles of chloroplast biology including electron transport (Iwai et al., 2010), photosynthetic regulation (Depège et al., 2003), assembly of photosynthetic complexes (Minai et al., 2006), and chloroplast genome segregation (Kobayashi et al., 2017). Thus, further study of the Chlamydomonas chloroplast is likely to continue to shed light on the chloroplast biology of land plants, including agriculturally important crop species.

**Figure 1.**
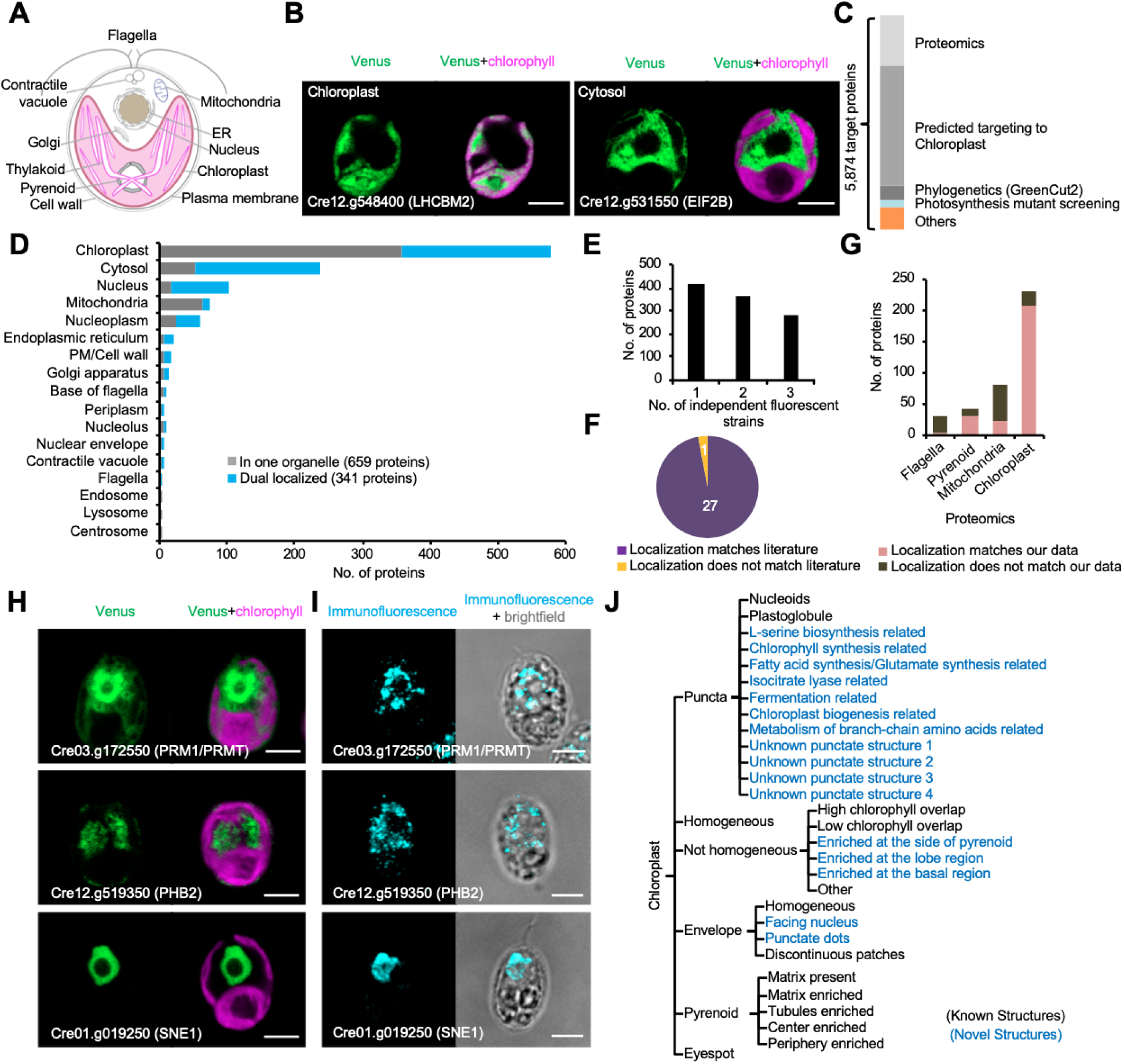
We assigned 1,032 tagged proteins to diverse localization patterns in 17 major compartments. (A) A diagram shows the cell structure of *Chlamydomonas reinhardtii*. (B) Representative images of the Venus-tagged chloroplast protein Cre12.g548400 (LHCBM2) and cytosol protein Cre12.g531550 (EIF2B). (C) A summary of target genes from various sources. (D) The number of proteins per subcellular location is shown, including proteins observed in one organelle (grey) and proteins observed in multiple organelles (blue). (E) The number of independent strains imaged for determining the localization patterns. (F) Agreement of our data with localizations identified in previous literature. (G) Comparison of our localizations with several proteomes in Chlamydomonas. (H) Our data are shown for proteins whose localizations disagree with proteomics-based localizations in Chlamydomonas. Cre03.g172550 (PRM/PRMT1) was previously found in the chloroplast proteome, Cre12.g519350 (PHB2) was previously found in the mitochondria proteome, and Cre01.g019250 (SNE1) was previously found in the flagella proteome. (I) Localizations of PRM1/PRMT, PHB2, and SNE1 in wild-type were assessed using immunofluorescence. (J) A decision tree for assigning chloroplast proteins to specific subcellular locations. Novel structures are labeled in blue. All scale bars are 5 μm.

In this study, we establish a comprehensive atlas of the subcellular localizations of 1,032 chloroplast candidate proteins in Chlamydomonas (Figure 1). Our results reveal novel chloroplast structures and spatial organization, new components of known cellular structures, and widespread dual localization of proteins. We also use this dataset to train a more accurate Chlamydomonas protein localization predictor. These insights and the associated plasmid, strain, and protein localization prediction resources open doors to the characterization of novel proteins and spatial organization of the chloroplast in green algae and land plants.

## RESULTS

### Establishing a Framework for Mapping Chloroplast Protein Localization

In order to maximize the total number of proteins localized in or associated with the chloroplast, we selected target proteins for fluorescent tagging from seven sources (Figure 1C and S1A; Table S1): 1) proteins homologous to those identified in the *Arabidopsis thaliana* chloroplast mass-spectrometry-based proteome (Ferro et al., 2010); 2) the Chlamydomonas mass-spectrometry-based chloroplast proteome (Terashima et al., 2010); 3) the Chlamydomonas mass-spectrometry-based pyrenoid proteome (Zhan et al., 2018); 4) proteins predicted to localize to the chloroplast by the bioinformatics software PredAlgo (Tardif et al., 2012); 5) GreenCut2 proteins (Karpowicz et al., 2011), which are a set of proteins conserved exclusively in the green lineage; 6) proteins identified to be required for photosynthesis based on Chlamydomonas mutant phenotypes (Li et al., 2019); and 7) other proteins we identified as potentially associated with chloroplast functions based on protein domains or annotations (see STAR Methods). To facilitate the classification of localizations, we included 29 proteins with known localization to the chloroplast or to other organelles in Chlamydomonas (Figure S1B).

To determine protein localizations, we used a previously established system (Mackinder et al., 2017) for expressing proteins tagged with the fluorescent protein Venus (see STAR Methods). Specifically, we cloned the open reading frame of each gene into a vector that drives expression from a constitutive promoter and appends a C-terminal fluorescent Venus tag (Nagai et al., 2002) for localization and three copies of the FLAG epitope (Hopp et al., 1988) for affinity purification of the tagged protein. We electroporated each construct into wild-type Chlamydomonas cells, which produced stable insertions at random sites within the genome (Zhang et al., 2014). We imaged the protein localizations in photoheterotrophically-grown live cells by using confocal fluorescence microscopy. While localizations can be affected by protein tags and non-native expression, we previously showed that such artifacts are rare with this pipeline (Mackinder et al., 2017).

### Localization Dataset Validation

We were successful in mapping the localization of 1,032 tagged proteins to 141 distinct localization patterns across 17 major organelles/cellular sites (Figure 1D; Table S2 and S3). To minimize experimenter bias in classifying the localization patterns, each localization image was independently analyzed by two researchers. The organelles/cellular site with the most localized proteins were the chloroplast (580 proteins), followed by the cytosol (238 proteins) (Figure 1D). These numbers include dual-localized proteins, which we will discuss later.

Next we evaluated the reliability of our data. Given that fluorescent tagging could lead to inaccurate protein localization due to either protein complex disruption or alteration of native regulation, we first investigated the reproducibility and agreement of our results with protein localizations from previous studies. Of the localized proteins, 62% were represented by at least two independent strains (Figure 1E). The localizations observed in the independent strains for a given protein agreed in >99% of the cases (Figure S1C and S1D). Furthermore, our localizations matched previously published localizations for 27 of the 28 Chlamydomonas proteins (96%) (Figure 1F; Table S4). The only exception was EZY1 (Cre06.g255750), which is normally expressed exclusively in early diploid zygotes and participates in the uniparental inheritance of the chloroplast genome (Armbrust et al., 1993). Since cells imaged in this study were haploid cells, the mis-localization of EZY1 to mitochondria in our data (Table S2), could be attributed to expression under non-native conditions, where the appropriate chloroplast targeting machinery may not be available.

We also analyzed the localization enrichment of a set of previously well-characterized proteins. All 32 known photosynthetic complex proteins and all 23 plastid ribosome proteins represented in our dataset were enriched in the chloroplast (Figure S1E). Taken together, the excellent agreement of our localization data with previous studies suggests that our dataset provides reliable localizations for uncharacterized proteins.

Our fluorescence images are particularly effective in validating reported organelle proteomics data and identifying potential contaminant proteins in those datasets (Figure 1G; Table S2). Our localization data suggest that 26 out of the 233 proteins from the published Chlamydomonas chloroplast proteome (Terashima et al., 2010) that are also represented in our dataset are actually not in the chloroplast under our experimental conditions. Similarly, 10 out of the 41 proteins previously detected in the pyrenoid proteome (Zhan et al., 2018), 56 out of the 81 proteins previously detected in the Chlamydomonas mitochondrial proteome (Atteia et al., 2009), and 21 out of the 25 reported high confidence flagellar proteome proteins (Pazour et al., 2005) do not match our localization data. We note that these numbers should not be interpreted as reflecting the overall accuracy of the mitochondrial or flagellar proteomes: we only tagged the subsets of these proteomes for which other omics evidence suggested a possible chloroplast localization, which enriches our dataset for mitochondrial or flagellar proteome false-positives.

To validate our data in these cases, we investigated the localizations of three proteins using antibodies to the native proteins by indirect immunofluorescence. The conserved histone-arginine N-methyltransferase (PRM1/PRMT: Cre03.172550) (Scebba et al., 2007), which had been previously detected in the chloroplast proteome (Terashima et al., 2010), was localized to the ER/nucleus in our data (Figure 1H). Prohibitin 2 (PHB2: Cre12.g519350) (Wang et al., 2010), which had been found in the mitochondrial proteome (Atteia et al., 2009), was localized to the cytosol (Figure 1H). A GreenCut2 protein (SNE1: Cre01.g019250) (Major et al., 2005), which had been previously detected in the flagellar proteome (Pazour et al., 2005), was localized to nucleoplasm (Figure 1H). Consistent with our localization dataset, we detected the native PRM1, PHB2, and SNE1 mainly in the ER/nucleus, cytosol, and nucleoplasm, respectively, by immunofluorescence (Figure 1I).

### Protein Localization Reveals Eleven Novel Chloroplast Punctate Structures Suggestive of Compartmentalized Biosynthetic Reactions

We assigned the 580 chloroplast proteins we characterized to one or more of 30 sub-chloroplast locations (Figure 1J). Among the most striking were 11 unique punctate localization patterns that we could not associate with previously described structures within the chloroplast (Figure 2 and S2; Table S2). The localization patterns differed in the number, diameter, and position of puncta within the chloroplast, suggesting that they correspond to distinct structures (Figure 2A, 2B, and 2C). We named the 7 previously-unnamed punctate-localized proteins <u>c</u>hloroplast <u>p</u>unctate <u>p</u>roteins (CPP1-7).

**Figure 2.**
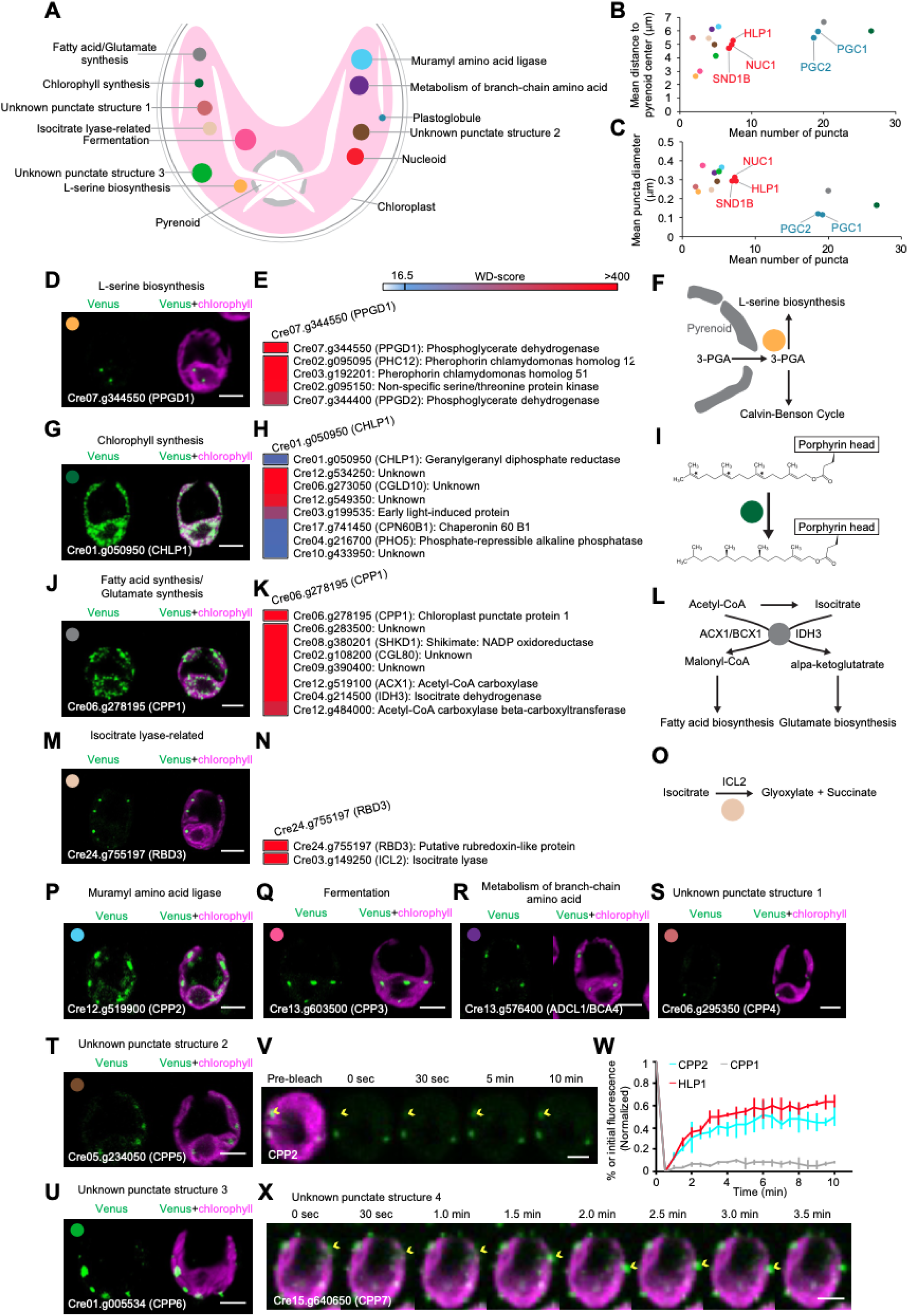
The data revealed novel punctate structures in the chloroplast. (A) A diagram illustrates the 12 chloroplast puncta structures we observed, 10 of which are novel. The relative size and approximate distance from the center of the pyrenoid are represented; for simplicity, only one of each structure is shown. Each structure is assigned a unique color that is used throughout this figure and Figure S2. (B-C) Novel punctate structures showed differences in the average position, number, and size of puncta. For each punctate structure, its mean distance to the pyrenoid center, the average number of puncta, and the mean punctum size are shown. Mean values ± SD from five independent cells are shown in Figure S2. (D) Representative images of Cre07.g344550 (PPGD1). (E) High-confidence interacting proteins of PPGD1. WD-scores represent our confidence in the interactions; scores greater than 16.5 correspond to the 3.7% highest-confidence interactions (see STAR Methods). (F) A diagram illustrates how the 3-phosphoglycerate dehydrogenase PPGD1 enhance its activity by localizing adjacent to the pyrenoid (G) Representative images of Cre01.g050950 (CHLP1). (H) High-confidence interacting proteins of CHLP1. (I) A diagram illustrates how the geranylgeranyl diphosphate reductase CHLP1 catalyzes the last step of the biogenesis of chlorophyll. (J) Representative images of Cre06.g278195 (CPP1). (K) High-confidence interacting proteins of CPP1. (L) A diagram illustrates how the CPP1 regulates the branching of metabolism between fatty acid synthesis and production of glutamate. (M) Representative images of Cre24.g755197 (RBD3). (N) High-confidence interacting proteins of RBD3. (O) A diagram shows how the RBD3 supports the isocitrate lyase ICL2. (P-U) Representative images of Cre12.g519900 (CPP2) (P), Cre13.g603500 (CPP3) (Q), Cre13.g576400 (ADCL1/BCA4) (R), Cre06.g295350 (CPP4) (S), Cre05.g234050 (CPP5) (T), Cre01.g005534 (CPP6) (U). (V) Fluorescence recovery of an CPP2-Venus punctum during 10 minutes after photobleaching of the whole punctum. The yellow arrow indicates the bleached punctum. (W) Fluorescence recovery profile of puncta of CPP2, Cre06.g285401 (HLP1),, and CPP1. Shown are the mean ± SD of 3 different puncta for each protein. (X) Representative images of Cre15.g640650 (CPP7) showing rapid movement of puncta. The yellow arrows indicate one of the puncta rapidly moving along the lobe of chloroplast. All scale bars are 5 μm.

To further explore these 11 novel structures and to identify additional components of each one, we performed immunoprecipitation-mass spectrometry on the tagged proteins. We identified on average 5 high-confidence protein interactors per structure, for a total of 59 proteins associated with these novel chloroplast punctate structures (Figure 2E, 2H, 2K, and 2N; Table S5). Many of the constituent proteins are conserved in land plants, suggesting the possibility that at least some of these structures are broadly conserved.

A number of the tagged proteins or their interactors correspond to metabolic enzymes, suggesting that these punctate structures may play functional roles in the spatial organization of biosynthetic reactions. Two themes emerge from the data: 1) Typically, only some of the enzymes of a pathway are localized to puncta, suggesting that the puncta enhance or regulate a subset of the reactions. 2) In some cases, punctate localization of an enzyme may allow it to perform its reaction at a location where its substrate is most available. These observations are consistent with previous observations of metabolism associated with cellular condensates (Castellana et al., 2014; Küken et al., 2018; O’Connell et al., 2012; Pareek et al., 2021). Below, we illustrate what we have been able to glean about the composition and potential functions of some of these novel structures:

### L-serine Biosynthesis

The conserved predicted 3-phosphoglycerate dehydrogenase Cre07.g344550 (Figure S3A), which catalyzes the commitment step of L-serine biosynthesis, localized to puncta, most of which were directly adjacent to the pyrenoid (Figure 2D). Since the pyrenoid is the site of production of 3-phosphoglycerate (3-PGA) by ribulose 1,5-bisphosphate carboxylase/oxygenase (Rubisco), the localization of 3-phosphoglycerate dehydrogenase to puncta next to the pyrenoid may enhance its activity through metabolic channeling. The pyrenoid is surrounded by presumably impermeable starch plates that are only punctured in a few places by thylakoid membranes (Engel et al., 2015); we speculate that the 3-phosphoglycerate dehydrogenase puncta localize to these openings to capture exiting 3-phosphoglycerate (Figure 2F). Cre07.g344550 co-precipitated with another predicted 3-phosphoglycerate dehydrogenase encoded adjacent to it in the genome, Cre07.g344400 (Figure 2E), suggesting that both enzymes may function in these puncta. Based on these observations, we propose to name these enzymes <u>P</u>yrenoid-associated 3-<u>P</u>hospho<u>g</u>lycerate <u>D</u>ehydrogenase PPGD1 and PPGD2, respectively, and the puncta glydehydrosomes.

### Chlorophyll Biosynthesis

CHLP1 (Cre01.g050950), a conserved predicted geranylgeranyl diphosphate reductase (Figure S3B) that catalyzes a series of reductions during the last step of the biogenesis of the key photosynthetic pigment chlorophyll (Tanaka et al., 1999), formed the most numerous puncta of all the novel structures we observed (Figure 2G). We did not observe a punctate localization for any of the 8 other enzymes in the chlorophyll biosynthesis pathway that we tagged and examined in our dataset (Table S2), suggesting that this is the only step of chlorophyll biosynthesis that benefits from being performed in puncta. Since CHLP1 needs to perform three separate reductions on its substrate (Figure 2I), we speculate that localization of CHLP1 to puncta increases the enzyme’s local concentration, allowing released product to more efficiently re-bind a CHLP1 active site during these sequential reductions.

CHLP1 physically interacted with Cre03.g199535, a conserved early light-induced protein with a chlorophyll *a*/*b* binding protein domain (Tanaka et al., 2010), and with the conserved but poorly-characterized protein CGLD10 (Cre06.g273050) (Figure 2H). These observations suggest that Cre03.g199535 and CGLD10 also play roles in chlorophyll biosynthesis, possibly by enhancing CHLP1’s function.

### Metabolic Regulation

The punctate-localized conserved protein Cre06.g278195, which we named CPP1 (Figure S3C), co-precipitated with two subunits of acetyl-CoA carboxylase, ACX1 (Cre12.g519100) and BCX1 (Cre12.g484000), and with the chloroplastic isocitrate dehydrogenase IDH3 (Cre04.g214500) (Figure 2J and 2K). Both of these enzymes perform essentially irreversible reactions downstream of citrate/acetyl-CoA. Thus, we hypothesize that the puncta formed by CPP1 regulate the branching of metabolism between fatty acid synthesis, which is downstream of acetyl-CoA carboxylase (Sasaki and Nagano, 2004), and the production of glutamate, which is downstream of isocitrate dehydrogenase (Elias and Givan, 1977) (Figure 2L).

### Glyoxylate Cycle

The conserved punctate-localized protein RBD3 (Cre24.g755197) (Figure 2M) contains a predicted Rubredoxin-like domain (Figure S3D), suggesting that it plays a role in electron transfer. RBD3 physically interacted with the predicted isocitrate lyase ICL2 (Cre03.g149250) (Figure 2N), a key enzyme in the glyoxylate cycle, which allows cells to metabolize two-carbon compounds such as acetate when simple sugars are not available (Figure 2O). Intriguingly, the glyoxylate cycle is thought to occur in peroxisomes (Kong et al., 2017) and was not previously thought to occur in the chloroplast. However, from our data and the literature, there is evidence suggesting that the enzymes necessary for the cycle are also present in the chloroplast. Specifically, we observed chloroplast localization of a malate dehydrogenase MDH1 (Cre03.194850) and a citrate synthase (Cre13.g579050) (Table S2). In addition, succinate dehydrogenase activity has been observed in spinach chloroplasts (Willeford et al., 1989), and bioinformatics (Table S7) (Tardif et al., 2012) predicts the chloroplast targeting of aconitases (Cre06.g252650 and Cre01.g004500), succinate dehydrogenase (Cre12.g528450), and fumarase (Cre06.g272500). Our observations therefore help support the possibility that chloroplasts are able to operate a glyoxylate cycle, which could increase the cell’s metabolic flexibility, and suggest that a portion of this cycle occurs in punctate structures.

### Punctate Structures Differ in their Exchange and Movement Dynamics

Intriguingly, the novel punctate structures we observe exhibit different dynamics in terms of the exchange of components with the chloroplast stroma and the movement of the structures within the chloroplast. Puncta of the predicted muramyl amino acid ligase CPP2 (Cre12.g519900) demonstrated rapid exchange of components with the stroma similar to the behavior of puncta of the chloroplast DNA-binding nucleoid component HLP1, as examined by fluorescence recovery after photobleaching (Figure 2V and 2W). In contrast, the punctate structure formed by CPP1, which we associated with branching of metabolism between fatty acid synthesis and the production of glutamate (Figure 2W), did not exhibit such rapid exchange. Moreover, whereas most structures did not move significantly during a 10-minute image acquisition, puncta that contained CPP7 (Cre15.g640650) showed rapid movement on the timescale of minutes (Figure 2X and Movie S1). We speculate that the rapid exchange of CPP2 and nucleoid components with stroma, and the rapid movement of CPP7 are important to the function of these compartments.

### Localization Data Reveals Novel Components of Chloroplast Substructures

In addition to discovering novel structures, we identified novel components across different known substructures within the chloroplast.

### Novel Nucleoid Components

One of two previously known punctate structures in our dataset were nucleoids (Figure 3A) (Kobayashi et al., 2017). Our dataset revealed two novel nucleoid proteins (Figure 3B), both of which co-precipitated and colocalized with the previously-characterized nucleoid protein HLP1 (Cre06.g285401) (Karcher et al., 2009) (Figure 3C and 3D). One of these novel proteins, Cre16.g672300, which we called <u>Nuc</u>leoid protein 1 (NUC1), is conserved in land plants and contains two predicted high mobility group protein domains (Figure S3E) that bind DNA (Mallik et al., 2018). The other protein, SND1B (Cre06.g256850), contains a predicted histone-lysine N-methyltransferase and a SAND DNA-binding domain (Bottomley et al., 2001). We hypothesize that these proteins mediate nucleoid function.

**Figure 3.**
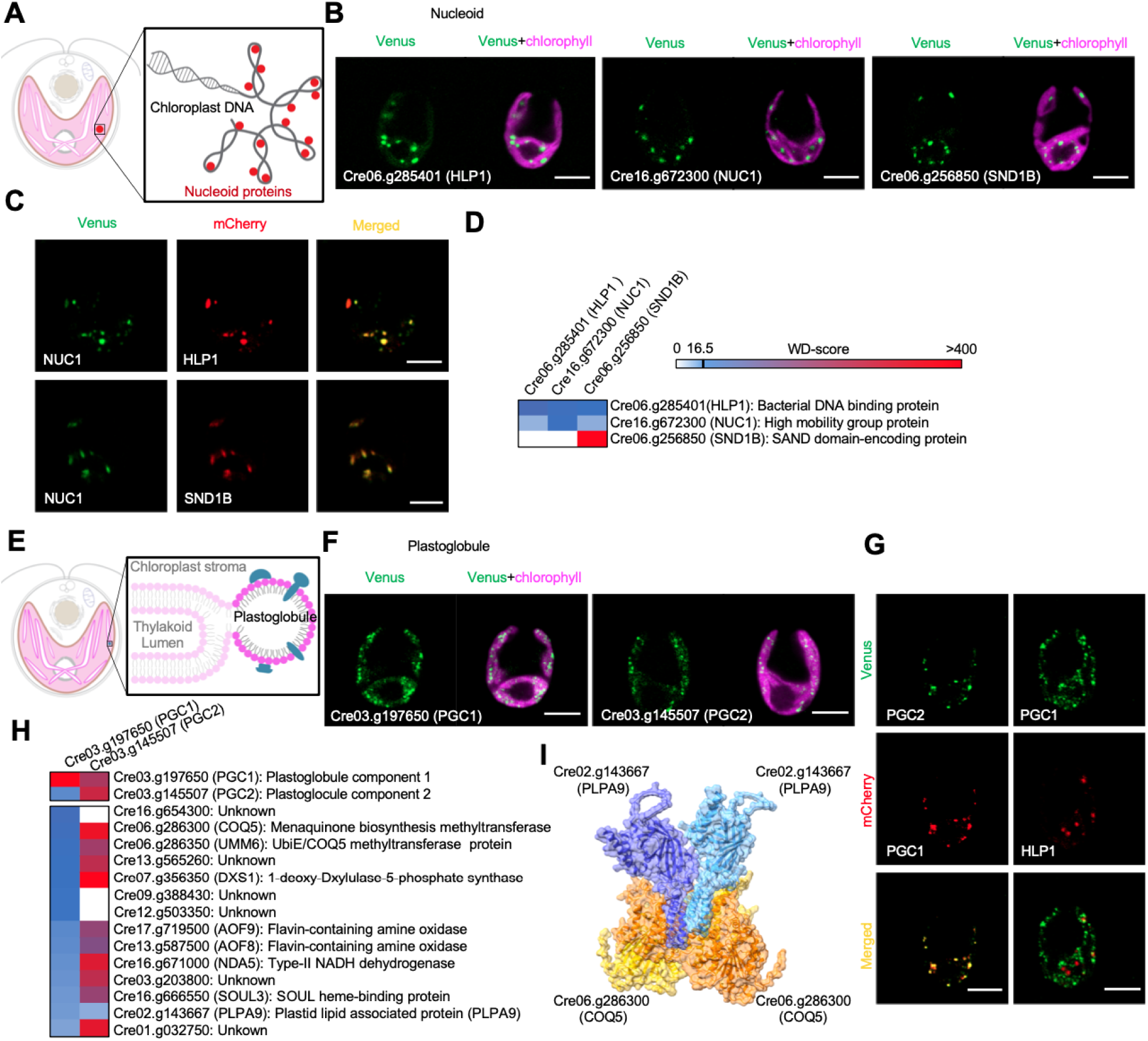
The data revealed novel components of nucleoids and plastoglobules in the chloroplast. (A) A diagram of the chloroplast nucleoid, in which the chloroplast DNA is organized into DNA-protein conglomerates. (B) Representative images of fluorescently-tagged Cre06.g285401 (HLP1), Cre16.g672300 (NUC1), and Cre06.g256850 (SND1B). (C) Dual tagging indicates the co-localization of HLP1, NUC1, and SND1B. (D) Protein-protein interactions among HLP1, NUC1, and SND1B. (E) Diagram of a plastoglobule, a thylakoid membrane-associated lipid droplet. (F) Representative images of fluorescently-tagged Cre03.g197650 (PGC1) and Cre03.g145507 (PGC2). (G) Dual tagging indicates the co-localization of PGC1 and PGC2. PGC1 was not co-localized with HLP1, indicating that they localize to distinct structures. (H) High-confidence interacting proteins of PGC1 and PGC2. (I) Structure modeling of Cre02.g143667 (PLPA9) and Cre06.g286300 (COQ5) by Alphafold suggests a direct interaction between these two proteins. All scale bars are 5 μm.

### Novel Plastoglobule Components

The other previously-known site of protein localization corresponded to plastoglobules, which are thylakoid membrane-associated lipid droplets containing triacylglycerols, plastoquinone, phylloquinone, carotenoids, and proteins related to their biosynthesis (Figure 3E) (Ytterberg et al., 2006). Our data revealed two novel plastoglobule-localized proteins, Cre03.g197650 and Cre03.g145507, which we named <u>p</u>lasto<u>g</u>lobule <u>c</u>omponent 1 and 2 (PGC1 and PGC2). PGC1 contains a PAP fibrilin domain found in structural proteins of plastoglobules (Deruere et al., 1994), which led us to hypothesize that the puncta it formed (Figure 3F) corresponded to plastoglobules. PGC2 showed very similar localization pattern to PGC1 (Figure 3F), co-localized with it (Figure 3G), and coprecipitated with it (Figure 3H), suggesting that they are part of the same structure. Immunoprecipitation of these two proteins pulled down six proteins whose homologs were previously found in the *Arabidopsis* plastoglobule proteome (Figure 3H) (Lundquist et al., 2012; Ytterberg et al., 2006). These proteins included the electron transport protein NAD5 (Cre16.g671000), the SOUL heme-binding protein SOUL3 (Cre16.g666550) (Shanmugabalaji et al., 2020), two phylloquinone biosynthesis related proteins UMM6 (Cre06.g286350) and COQ5 (Cre06.g286300) (Gross et al., 2006; Lee et al., 1997), and the plastid lipid-associated protein PLPA9, which we computationally predict forms a complex with COQ5 (Figure 3I). Based on this, we conclude that PGC1 and PGC2 are novel plastoglobule proteins.

Interestingly, our immunoprecipitation experiments also identified enzyme involved in processes not previously thought to occur at the plastoglobules. Specifically, we found DXS1 (Cre07.g356350), a conserved protein predicted to be a synthase for 1-deoxy-D-xylulose 5-phosphate (Figure S3F), which generates a precursor for isoprenoid and vitamin B_1_ and B_6_ synthesis (Lois et al., 1998). We also found in the immunoprecipitation AOF8 (Cre13.g587500) and AOF9 (Cre17.g719500), two conserved predicted flavin-containing amine oxidases that catalyze the oxidative cleavage of alkylamines into aldehydes and ammonia. These findings suggest that plastoglobules perform previously unappreciated functions in the metabolism of 1-deoxy-D-xylulose 5-phosphate and alkylamines.

### Novel Pyrenoid Components

The pyrenoid is a non-membrane-bound proteinaceous sub-organelle of the chloroplast in which the rate of CO_2_ fixation into organic carbon is enhanced by supplying the CO_2_-fixing enzyme Rubisco with a high concentration of CO_2_ (Figure 4A) (Osafune et al., 1990; Wunder et al., 2019). Within our dataset, we observed the localization of 18 novel proteins localized to the pyrenoid periphery, matrix, tubules, or pyrenoid center (Table S2). Two of the pyrenoid matrix-localized proteins, the predicted histone deacetylase HDA5 (Cre06.g290400) and uncharacterized protein Cre16.g648400, harbor predicted Rubisco-binding motifs (Meyer et al., 2020), suggesting that they bind directly to Rubisco (Figure 4B).

**Figure 4.**
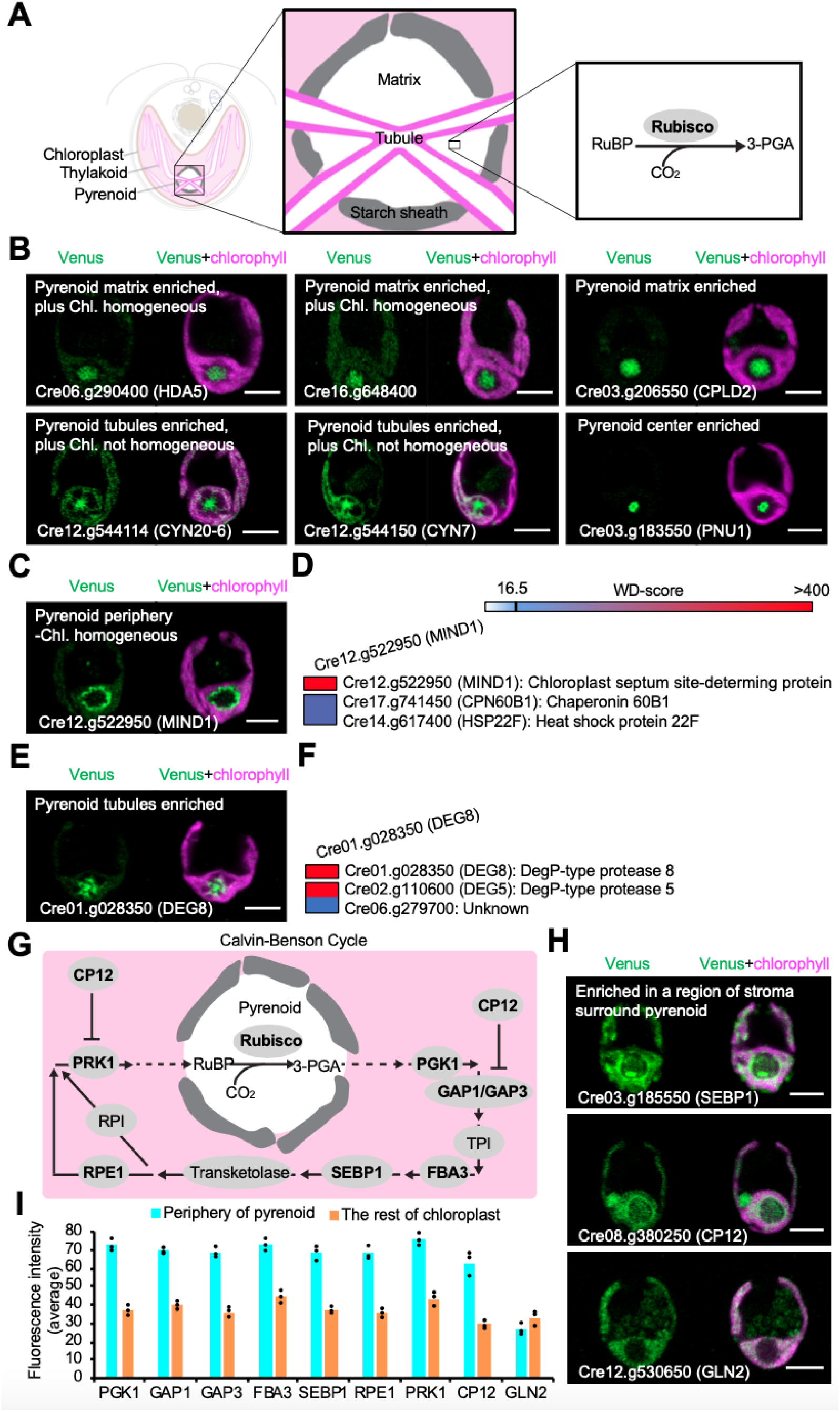
Our data revealed novel pyrenoid components and the enrichment of Calvin-Benson cycle enzymes in the stroma around the pyrenoid. (A) A diagram of the pyrenoid highlights the starch sheath, pyrenoid tubules, and pyrenoid matrix where most of the carbon-fixing enzyme Rubisco is located. (B) Representative images of Cre06.g290400, Cre16.g648400, and Cre03.g206550 (CPLD2), enriched at the pyrenoid matrix; Cre12.g544114 (CYN20-6) and Cre12.g544150 (CYN7) enriched in the pyrenoid tubules; and Cre03.g183550 (PNU1) localized to the pyrenoid center. (C) Representative images of Cre12.g522950 (MIND1) enriched at the pyrenoid periphery. (D) High-confidence interacting proteins of MIND1. (E) Representative images of Cre01.g028350 (DEG8) enriched in the pyrenoid tubules. (F) High-confidence interacting proteins of DEG8. (G) Simplified model of the Calvin-Benson cycle in Chlamydomonas. (H) Representative images of Cre03.g185550 (SEBP1) and Cre08.g380250 (CP12) enriched in a region of the stroma immediately surrounding the pyrenoid. For comparison, Cre12.g530650 (GLN2, Glutamine synthetase), homogeneously localized throughout the stroma. (I) Calvin-Benson cycle enzymes showed 1.85-fold enrichment in the stroma around pyrenoid. The measurement of fluorescence intensity was conducted with Fiji (ImageJ). The measured value and mean from three independent cells are shown. All scale bars are 5 μm.

MIND1 (Cre12.g522950), the Chlamydomonas homolog of the Arabidopsis chloroplast division site regulator MinD1 (Fujiwara et al., 2017), was enriched at the pyrenoid periphery (Figure 4C). MIND1 co-precipitated with plastid chaperonin 60 beta 1 subunit CPN60B1 (Cre17.g741450) (Figure 4D), whose Arabidopsis homolog has also been implicated in plastid division (Suzuki et al., 2009), suggesting the conservation of this interaction in plastid division in algae. Considering that the pyrenoid typically divides by fission during chloroplast division (Freeman Rosenzweig et al., 2017), we hypothesize that MIND1’s localization to the pyrenoid periphery plays a role in coordinating pyrenoid fission with chloroplast division.

The predicted phosphoglycolate phosphatase CPLD2 (Cre03.g206550) (Schwarte and Bauwe, 2007) was enriched in the pyrenoid matrix (Figure 4B). Phosphoglycolate phosphatase consumes 2-phosphoglycolate, a competitive inhibitor of the Calvin-Benson cycle enzymes triose phosphate isomerase (Wolfenden, 1969) and sedoheptulose 1,7-bisphosphate phosphatase (Flügel et al., 2017). The localization of CPLD2 to the pyrenoid likely allows the cell to consume 2-phosphoglycolate at its source near Rubisco before it can exit the pyrenoid and inhibit the activity of key enzymes in the surrounding chloroplast stroma.

The pyrenoid tubules are modified thylakoid membranes that traverse the pyrenoid and are thought to supply it with concentrated CO_2_. We observed 9 proteins localizing to the pyrenoid tubules, including two predicted peptidyl-prolyl cis-trans isomerases (CYN20: Cre12.g544114 and CYN7: Cre12.g544150) (Figure 4B) and the predicted DegP-type protease DEG8 (Cre01.g028350) (Figure 4E), which co-precipitated with another DegP-type protease, DEG5 (Figure 4F). These observations suggest that tubules may have a role in protein folding, degradation, and/or import of new proteins into the pyrenoid.

Finally, our data support a role for the pyrenoid in nucleic acid degradation in algae. The bifunctional nuclease domain-containing protein Cre03.g183550, which we named <u>p</u>yrenoid <u>nu</u>clease 1 (PNU1), localized to the pyrenoid center (Figure 4B). In plants, bifunctional nucleases are responsible for the degradation of RNA and single-stranded DNA in several biological processes (Pérez-Amador et al., 2000). Considering that oxidized RNA localizes to the pyrenoid in Chlamydomonas (Zhan et al., 2015), we speculate that the pyrenoid may be a site of degradation of oxidized RNA. Localizing RNA-degrading enzymes to the pyrenoid could allow for increased specificity of degradation for damaged RNA.

### Calvin Cycle Enzymes are Enriched in the Stroma Surrounding the Pyrenoid

The Calvin Cycle is the metabolic cycle that enables the assimilation of CO_2_. It consists of the CO_2_-fixing enzyme Rubisco and 11 other enzymes that convert Rubisco’s product, phosphoglycerate, into its substrate, ribulose-1,5-bisphosphate, allowing the cycle to continue. In Chlamydomonas and likely in all pyrenoid-containing algae, Rubisco is the only Calvin Cycle enzyme present in the pyrenoid, while the other enzymes are all in the stroma (Figure 4G) (Küken et al., 2018).

From our dataset, we observed that the Calvin Cycle enzyme sedoheptulose-1,7-bisphosphatase SEBP1 and the Calvin Cycle regulatory protein CP12 were both enriched in a region of the stroma immediately surrounding the pyrenoid (Figure 4H). This enrichment around the pyrenoid had not been noticed in our previous study (Küken et al., 2018) because of the lack of other stromal-localized proteins for comparison. Revisiting those data and reexamining the localization of the proteins again under the current growth conditions, it is now apparent that the Calvin Cycle enzymes phosphoglycerate kinase (PGK1), glyceraldehyde 3-phosphate dehydrogenase (GAP1 and GAP3), fructose-1,6-bisphosphate aldolase (FBA3), sedoheptulose-1,7-bisphosphatase (SEBP1), ribulose phosphate-3-epimerase (RPE1), and phosphoribulokinase (PRK1) are all enriched in the region of the stroma immediately surrounding the pyrenoid (Figure 4I and S4). The enrichment of these enzymes in the periphery of the pyrenoid may enhance the activity of the Calvin Cycle, considering that Rubisco is resident inside the pyrenoid and its substrates and products must therefore diffuse in and out of the periphery of the pyrenoid. These observations motivate questions for future research, including: how are these enzymes localized to the periphery of the pyrenoid and how does their localization change under high CO_2_, where some of the Rubisco dissolves into the stroma?

### Data Reveal Unexpected Thylakoid Associations and Protein Distributions

Our data suggest the unexpected thylakoid association of several proteins, as well as reveal an intriguing gradient distribution for one thylakoid protein. The thylakoid membranes are the site where light energy is captured by chlorophyll pigments used to generate ATP and NADPH for the cell. Of the proteins with non-homogeneous chloroplast localization in our dataset, 40 exhibited high localization overlap with chlorophyll (Figure 5A and 5B), while 31 exhibited low overlap (Figure 5C). We interpret high localization overlap with chlorophyll as indicative of thylakoid membrane association. Indeed, of the 71 proteins with non-homogeneous localization patterns, all 11 proteins with transmembrane domains showed high chlorophyll overlap (p=0.001, Fisher’s exact test). Below, we illustrate how our observation of thylakoid membrane association advances our understanding of protein functions.

**Figure 5.**
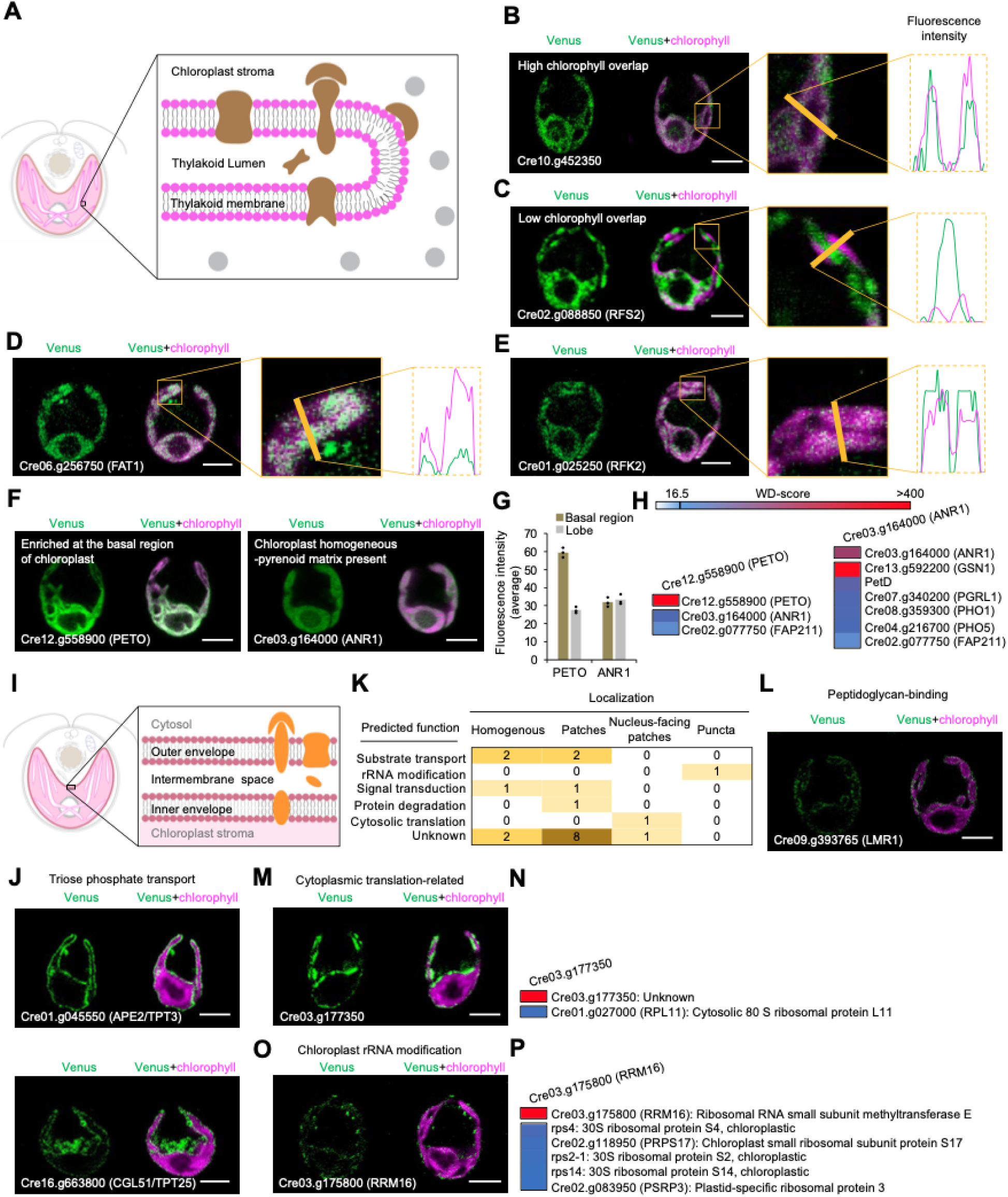
Our data revealed thylakoid-associated enzymes, the enrichment of PETO at the base of the chloroplast, and regions of the chloroplast envelope specialized for different functions. (A) A diagram shows the thylakoid membrane, thylakoid-associated enzymes (brown), and non-thylakoid-associated enzymes (grey). (B) Representative images and line intensity profile of fluorescently-tagged Cre10.g452350, which showed high overlap with chlorophyll. The measurement of fluorescence intensity was conducted with Fiji (ImageJ). (C) Representative images and line intensity profile of Cre02.g088850 (RFS2), which showed low overlap with chlorophyll. (D) Representative images and line intensity profile of Cre06.g256750 (FAT1), which showed high chlorophyll overlap. (E) Representative images and line intensity profile of Cre01.g025250 (RFK2), which showed high chlorophyll overlap. (F) Representative images of Cre12.g558900 (PETO), which was enriched in the base of the chloroplast. ANR1 (Cre03.g164000) homogeneously localized throughout the chloroplast. (G) PETO showed a 2-fold enrichment of average fluorescence intensity in basal region. The average fluorescence intensity in the basal or lobe region are shown for three independent cells; bars represent mean values. (H) High-confidence interacting proteins of PETO and ANR1. GSN1, glutamate synthase; PetD, cytochrome b6f subunit IV; PGRL1, proton-gradient related-like; PHO1 and PHO5, alkaline phosphatases. (I) A diagram of the chloroplast envelope highlights the outer membrane, intermembrane space, inner membrane, and chloroplast envelope-associated proteins. (J) Representative images of Cre01.g045550 (APE2/TPT3) and Cre16.g663800 (CGL51/TPT25), homogeneously localized throughout the chloroplast envelope. (K) A heat map shows the observed localization and predicted function of 20 chloroplast envelope proteins. (L) Representative images of Cre09.g393765 (LMR1), which localized to patches along the chloroplast envelope. (M) Representative images of Cre03.g177350, which localized to the nucleus-facing portion of the chloroplast envelope. (N) High-confidence interacting proteins of Cre03.g177350. (O) Representative images of Cre03.g175800 (RRM16), which localized as punctate dots along the chloroplast envelope. (P) High-confidence interacting proteins of Cre03.g175800 (RRM16). All scale bars, 5 μm.

In photosynthetic eukaryotes, fatty acids are made by fatty acid synthase in the chloroplast stroma (Walker and Harwood, 1985), but the localization of the enzymes that process the nascent fatty acids has not been completely defined. Interestingly, our data suggest that the only predicted chloroplastic acyl-ACP thioesterase FAT1 (Cre06.g256750), which releases fatty acids from fatty acid synthase (Hölzl and Dörmann, 2019), is associated with thylakoid membranes (Figure 5D). This observation suggests that nascent fatty acids are released in the proximity of thylakoid membranes, into which they may initially partition.

Our data also suggest that riboflavin kinase RFK2 (Cre01.g025250) is associated with the thylakoid membrane (Figure 5E). Riboflavin kinase phosphorylates riboflavin to produce flavin mononucleotide, an essential cofactor for a variety of enzymes including the thylakoid-localized NADH dehydrogenase (Tsibris et al., 1966). However, the localization of riboflavin kinase within the chloroplast was not previously known. The localization of RFK2 to the thylakoid membrane suggests that flavin mononucleotide is produced in proximity to where it is needed for assembly into NADH dehydrogenase.

In addition to the new proteins associated with the thylakoid, we also uncovered an intriguing distribution of a known thylakoid-associated protein, PETO (Cre12.g558900). PETO has been proposed to be important for photosynthetic cyclic electron flow, a poorly-understood pathway of photosynthesis that pumps additional protons across the thylakoid membrane without producing net reducing equivalents (Takahashi et al., 2016). In our dataset, PETO stood out as the only protein that showed a gradient localization pattern across the chloroplast, with a two-fold enrichment at the base of the chloroplast (Figure 5F, 5G, and S5A). While the specific function of PETO in cyclic electron flow remains unknown, our observation that it is localized in a gradient suggests the possibility that cyclic electron flow may be more active at the base of the chloroplast. This activity could result in pumping of additional protons into the thylakoid lumen in the proximity of the pyrenoid, where they are needed to drive the conversion of HCO_3_^-^ to CO_2_ by carbonic anhydrase (Pronina and Semenenko, 1990; Raven, 1997).

Our immunoprecipitation and mass spectrometry data examining interactors confirm the previously observed physical interaction of PETO with the cyclic electron flow regulator ANR1 (Cre03.g164000) (Takahashi et al., 2016) (Figure 5H). However, unlike PETO, ANR1 did not show a gradient localization (Figure 5F) and affinity purification of ANR1 did not yield detectable amounts of PETO (Figure 5H, Table S5), suggesting that only a fraction of ANR1 is associated with PETO. In addition, ANR1 co-precipitated with Cytochrome *b*_6_*f* subunit IV (PetD) and with the cyclic electron flow regulator PGRL1 (Cre07.g340200) (Terashima et al., 2012) (Figure 5H), supporting a possible direct role of ANR1 in the regulation of cyclic electron flow. The highest-confidence interactor of ANR1 was the predicted NADH-dependent glutamate synthase GSN1 (Cre13.g592200), suggesting the possibility that ANR1 could be downregulating cyclic electron flow in response to increased needs for NADPH by glutamate synthase.

### Chloroplast Envelope Localization Patterns Suggest Functionally Specialized Regions

The chloroplast envelope, as the interface between the chloroplast and surrounding cytosol, controls the exchange of ions, metabolites, proteins, and signals (Figure 5I). Surprisingly, out of the 20 chloroplast envelope-localized proteins, only five showed a homogeneous localization throughout the envelope (Figure 5J, 5K, and S5B), whereas all of the other 15 showed one of at least three distinct heterogeneous localization patterns: patches (12 proteins), nucleus-facing patches (2 proteins), and puncta (1 protein) (Figures 5K-M, 5O, and S5C-S5E; Table S2). It is possible that the patches represent multiple distinct structures, as the proteins with these localization patterns did not share any high-confidence protein interactions (Table S5). These observations suggest that most proteins operate in specialized regions at the chloroplast envelope. Below, we discuss protein functions associated with each localization pattern.

Proteins localized to patches along the chloroplast envelope included LMR1 (Cre09.g393765), which contains two predicted peptidoglycan-binding LysM domains (Mesnage et al., 2014) (Figure 5L). While some chloroplasts are surrounded by peptidoglycan, as in the moss *Physcomitrella patens* (Hirano et al., 2016), the apparent absence of most of the peptidoglycan biosynthesis genes in the Chlamydomonas genome suggest that LMR1 instead binds other glycans at the chloroplast envelope.

Proteins localized to nucleus-facing patches included the conserved protein Cre03.g177350 (Figure 5M and S3G). This protein physically interacted with the cytosolic 80S ribosomal protein L11 (Figure 5N), suggesting that Cre03.g177350 could be involved in the cytosolic translation of chloroplast proteins prior to their import into the chloroplast.

The protein that localizes to puncta along the chloroplast envelope (Figure 5O) is the conserved protein RRM16 (Cre03.g175800) (Figure S3H), which bears two ribosomal RNA (rRNA) methyltransferase domains. This localization suggests that the chloroplast envelope could also be a site where rRNA modification takes place. Consistent with this hypothesis, our mass spectrometry data showed high-confidence interactions between RRM16 and several chloroplast ribosome small subunit components, including rps4, PRPS17, rps2-1, rps14, and PSRP3 (Figure 5P). The presence of a predicted chloroplast targeting sequence in RRM16 (Tardif et al., 2012) and its physical interactions with chloroplast-encoded ribosome subunits suggest that RRM16 is acting on chloroplast rRNA rather than cytosolic rRNA. The localization of chloroplast rRNA modification to the chloroplast envelope could provide an opportunity for cytosolic signals to regulate the chloroplast ribosome.

### Many Proteins Have Unexpected Localizations to Multiple Compartments

Localization of a single gene’s protein products to multiple cellular compartments is a widespread phenomenon (Carrie and Whelan, 2013; Krupinska et al., 2020; Thul et al., 2017), which can enable signaling between organelles (Isemer et al., 2012a, 2012b) or increase the number of coding products within a restricted genome size (Carrie et al., 2009).

We identified 341 proteins with multiple compartment localization (Figure 6A), substantially more than the approximately 250 that have been identified across all previous studies in plants to date (Carrie and Whelan, 2013). We observed multiple targeting in 87 distinct localization patterns (Figure 6B and Table S3), six times more distinct patterns than seen previously in plants. Four proteins were multiply localized to 4 compartments (Figure 6A and 6C).

**Figure 6.**
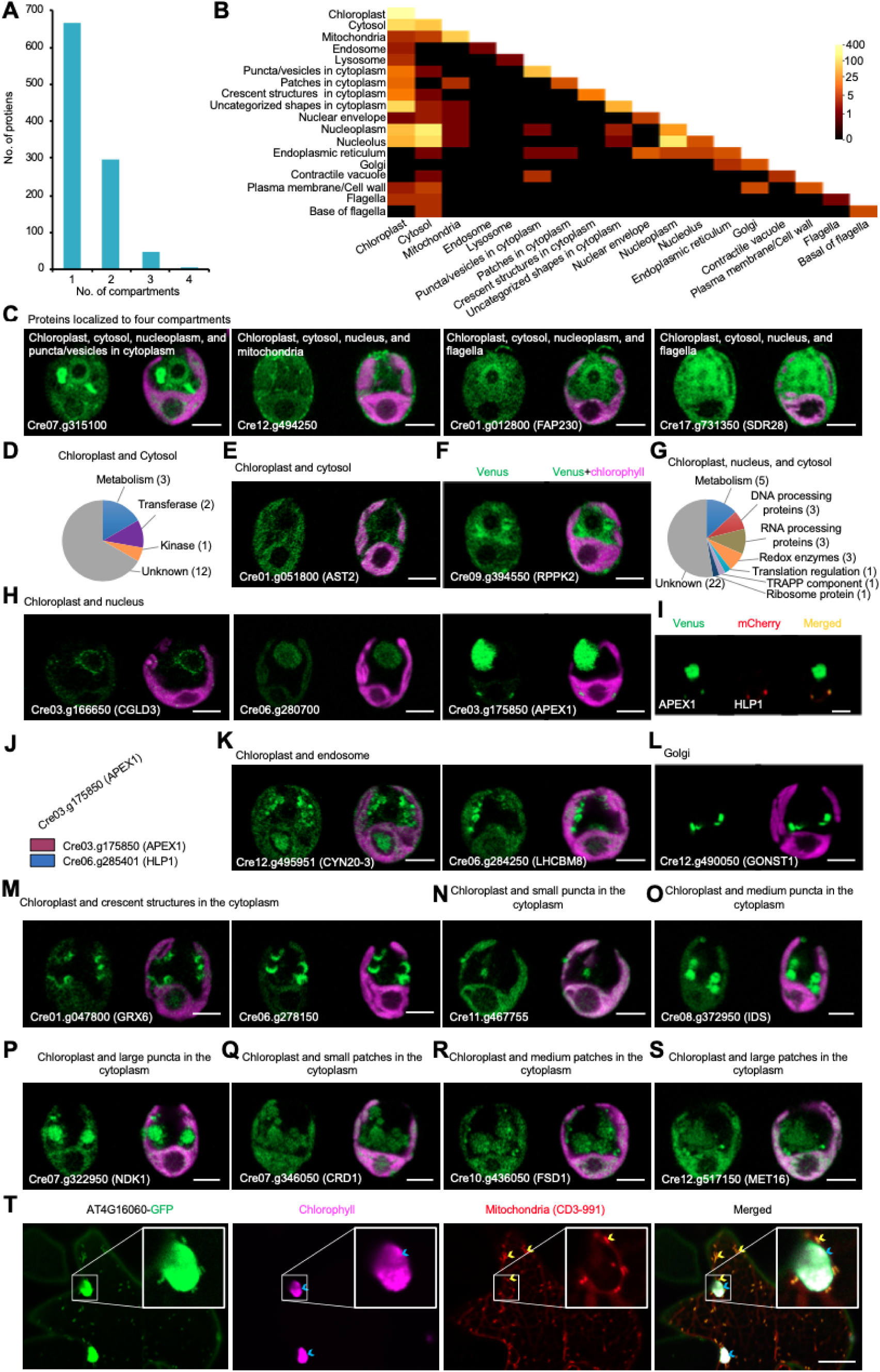
Many proteins localized to multiple compartments. (A) The number of proteins localized to 1, 2, 3, or 4 compartments is shown. (B) A heat map shows observed dual localizations. (C) Representative images of proteins localized to four compartments. (D) Functional classification of 18 proteins dual-localized to the chloroplast and cytosol. (E) Representative images of Cre01.g051800 (AST2), dual-localized to the chloroplast and cytosol. (F) Representative images of Cre09.g394550 (RPPK2), which localized to the chloroplast, cytosol, and nucleoplasm. (G) Functional classification of 39 proteins dual-localized to the chloroplast, nucleus, and cytosol. (H) Representative images of proteins dual-localized to the chloroplast and nucleus. (I) Dual tagging revealed the co-localization of Cre03.g175850 (APEX1, a putative exodeoxyribonuclease III), and HLP1 in the chloroplast. (J) The IP-MS data revealed a high-confidence interaction between APEX1 and HLP1. (K) Representative images of Cre12.g495951 (CYN20-3) and Cre06.g284250 (LHCBM8), dual-localized to the chloroplast and endosome. (L) Representative images of Cre12.g490050 (GONST1) whose homolog in Arabidopsis (AT2G13650) is localized to the Golgi apparatus. (M) Representative images of Cre01.g047800 (GRX6) and Cre06.g278150, dual-localized to the chloroplast and crescent structures in the cytoplasm. (N) Representative images of Cre11.g467755, dual-localized to the chloroplast and small puncta in the cytoplasm. (O) Representative images of Cre08.g372950 (IDS1), dual-localized to the chloroplast and medium puncta in the cytoplasm. (P) Representative images of Cre07.g322950 (NDK1), dual-localized to the chloroplast and large puncta in the cytoplasm. (Q) Representative images of Cre07.g346050 (CRD1), dual-localized to the chloroplast and small patches in cytoplasm. (R) Representative images of Cre10.g436050 (FSD1), dual-localized to the chloroplast and medium patches in cytoplasm. (S) Representative images of Cre12.g517150 (MET16), dual-localized to the chloroplast and large patches in cytoplasm. (T) Representative images of AT4G16060, the Arabidopsis homolog of Cre12.g494250, in tobacco leaf cells. AT4G16060 was observed in the chloroplast, cytosol, and mitochondria. The yellow arrows indicate mitochondria labeled by mitochondria mCherry marker CD3-991. The blue arrows indicate chloroplasts. All scale bars are 5 μm.

Because of how we selected our proteins for localization, our dataset is particularly enriched in proteins where one of the multiple sites of localizations is the chloroplast. Of the 341 multiple-localized proteins, 214 proteins were dual targeted to the chloroplast and one of 13 other regions, including the cytosol, mitochondria, endosome, lysosome, puncta/vesicles in the cytoplasm, patches in the cytoplasm, crescent structures in the cytoplasm, uncategorized shapes in the cytoplasm, the nuclear envelope, the nucleoplasm, the nucleolus, the plasma membrane/cell wall, and the flagella (Figure 6B and S6A-S6U; Table S2).

### Chloroplast and Cytosol

We observed 16 proteins with clear dual localizations to the chloroplast and cytosol (Figure 6D, 6E, and S6A). Many of these proteins contained predicted enzymatic domains (Figure 6D), suggesting that they are enzymes that function in both compartments. In some cases, our observed dual localizations identify candidate enzymes for activities that have been observed biochemically in those compartments. For example, the activity of ribose-phosphate pyrophosphokinase, which catalyzes a key step in purine nucleotide synthesis, has been detected in both the chloroplast and cytosol in spinach (Krath and Hove-Jensen, 1999), but the protein responsible for the activity in the chloroplast has not previously been identified. Our observation that the conserved ribose-phosphate pyrophosphokinase RPPK2 (Cre09.g394550) (Figure S3I) shows dual localization to both the cytosol and chloroplast suggests that this enzyme mediates the synthesis of phosphoribosyl diphosphate in both compartments (Figure 6F).

We also observed 31 proteins with a primary fluorescence signal in the chloroplast and relatively weak signal in the cytosol (Figure S6B). Some of these proteins are likely to be functional only in the chloroplast, as they are components of the photosynthetic apparatus or of the plastid ribosome (Table S2). The observation of these proteins in the cytosol may reflect a longer cytosolic residence time before their import into chloroplast (Jarvis and Robinson, 2004), or this could be an overexpression artifact of our system.

### Chloroplast and Nucleus

Our dual localization data suggest that the chloroplast and nucleus share nucleic acid processing and repair factors. We identified five proteins showing exclusively chloroplast and nucleus localizations (Figure 6G, 6H, and S6E). These proteins all had predicted functions related to nucleic acids. Of these five, the conserved predicted RNA helicase CGLD3 (Cre03.g166650) and putative RNA splicing factor Cre06.g280700 both localized to the nucleus and throughout the chloroplast (Figure 6H), suggesting that both proteins act on RNA in both compartments. The conserved DNA repair exonuclease APEX1 (Cre03.g175850) (Figure S3J) localized to both the nucleus and chloroplast nucleoids (Figure 6H), and co-localized and co-precipitated with the nucleoid component HLP1 (Figure 6I and 6J), suggesting that it contributes to the repair of both genomes.

### Chloroplast and Endosome or Lysosome

We observed 6 proteins localized to the chloroplast and either the endosome or the lysosome (Figure 6K, and S6K-S6M). For some of these proteins, this dual localization likely reflects a functional role of the protein in both compartments. For example, the conserved peptidyl-prolyl cis-trans isomerase CYN20-3 (Cre12.g495951) (Figure S3K) could isomerize prolines in both the chloroplast and in the endosome (Figure 6K). However, other proteins showing dual chloroplast and endosome localization, such as the light-harvesting protein LHCBM8 (Cre06.g284250) (Figure 6K), are likely proteins that function in the chloroplast and are degraded in the lysosome by chlorophagy (Ishida et al., 2008; Wolfe et al., 1997).

### Chloroplast and Crescent Structures in the Cytoplasm

Among the most striking dual localizations were proteins that localized to the chloroplast and crescent structures in the cytoplasm, a localization pattern that has not been described previously to our knowledge (Figure 6M). Depending on the localized protein, the crescent structures were either small (∼1 μm) or medium-sized (∼2 μm); these two sizes could represent either distinct structures or different stages of development of the same structure. The structures did not appear to be Golgi (Figure 6L), endosomes (Figure S1B and S6L), or lysosomes (Figure S6M), as the latter organelles showed different localizations and morphologies.

The predicted domains of 9 out of 20 proteins that localize to these crescent structures suggest that the crescent structures play roles in nucleotide and phosphate metabolism (Table S2). These proteins included the predicted polynucleotide phosphatase/kinase Cre11.g467709 (Figure 6V), the predicted purine biosynthesis enzyme Cre17.g734100 (Figure S6N), and the predicted phosphate transporter Cre07.g325740 (Figure S6V). Since cellular phosphate is primarily used for nucleotide biosynthesis, it makes sense that these two functions would be spatially co-localized.

The predicted functions, localization patterns, and size of the structures together suggest that the crescent localizations correspond to the matrix of acidocalcisomes, poorly-characterized vesicular structures that store phosphate as a single large spherical granule of polyphosphate (Aksoy et al., 2014; Docampo et al., 2005; Komine et al., 2000; Ruiz et al., 2001). Fluorescently-tagged proteins localizing to the matrix of acidocalcisomes would show a crescent structure due to their exclusion from the spherical polyphosphate granule (Figure 6M, S6N, and S6V). Indeed, the matrix of acidocalcisomes observed by electron microscopy (Aksoy et al., 2014) appeared as crescents of similar size to the structures we observed by microscopy.

Acidocalcisomes are possibly the only organelle conserved from bacteria to plants and humans (Docampo et al., 2005). They are essential for cellular survival under nutrient deprivation, but we are only beginning to understand their protein composition in any organism (Huang et al., 2014). Our identification of 20 candidate acidocalcisome proteins advances the molecular characterization of these fascinating structures. Moreover, the relatively large number (12) of proteins dual-localized to the chloroplast and these structures suggests that there could be extensive interactions between chloroplasts and acidocalcisomes, with potential for cycling of phosphate between the two compartments.

### Chloroplast and Other Structures

We observed 27 proteins dual-localized to the chloroplast and cytoplasmic puncta of one of three different diameters: small (∼1 μm), medium (∼2 μm), or large (∼3 μm) (Figure 6N-6P and S6O-S6Q). All three classes of puncta contained proteins with predicted enzymatic domains (Table S2), but contained no homologs of well-characterized proteins, precluding us from conclusively assigning these localizations to known structures. We also observed 12 proteins dual-localized to the chloroplast and small (∼2 μm diameter), medium (∼3.5 μm), or large (∼5 μm) cytoplasmic patches (Figure 6Q-6S and S6R-S6T), or to one of many other uncategorized shapes in the cytoplasm (Figure S6U). Future characterization of the functions and spatial dynamics of these proteins is likely to provide novel insights into the cell biology of plants and algae.

### Machine Learning Enables Proteome-Wide Protein Localization Predictions

Machine learning provides an opportunity to expand the scope of the protein localization findings of the present study to the genome-wide scale. The current state-of-the-art predictor for Chlamydomonas protein localization, PredAlgo (Tardif et al., 2012), has been a tremendously useful resource for the scientific community. However, it was trained on a relatively small dataset of 152 proteins. The much larger number of protein localizations resulting from this present work combined with advances in machine learning classification of protein sequences allowed us to train a more accurate protein localization predictor.

We built our new predictor, PB-Chlamy, based on ProtBertBFD (Elnaggar et al., 2021), a natural language processing model of protein features pre-trained on the BFD database (Steinegger et al., 2019) containing 2.5 billion protein sequences from diverse organisms. We trained three separate ProtBertBFD protein sequence classifiers on Chlamydomonas protein localization data (Figure 7A): one each to recognize chloroplast, mitochondrial, and secretory proteins. For each localization category we generated a combined dataset using the data in the present work, our previous protein localization study (Mackinder et al., 2017), and the training dataset assembled for PredAlgo (Tardif et al., 2012). Each dataset is composed of a set of positives, i.e. proteins known to localize to a particular subcellular location, and negatives, i.e. proteins found to not localize to the location. We split each dataset into training, validation, and testing subsets with a 3:1:1 ratio (Table S7). We used the training set to train a linear classifier to distinguish proteins that do or do not localize to a compartment, evaluating against the validation set of proteins during training.

**Figure 7.**
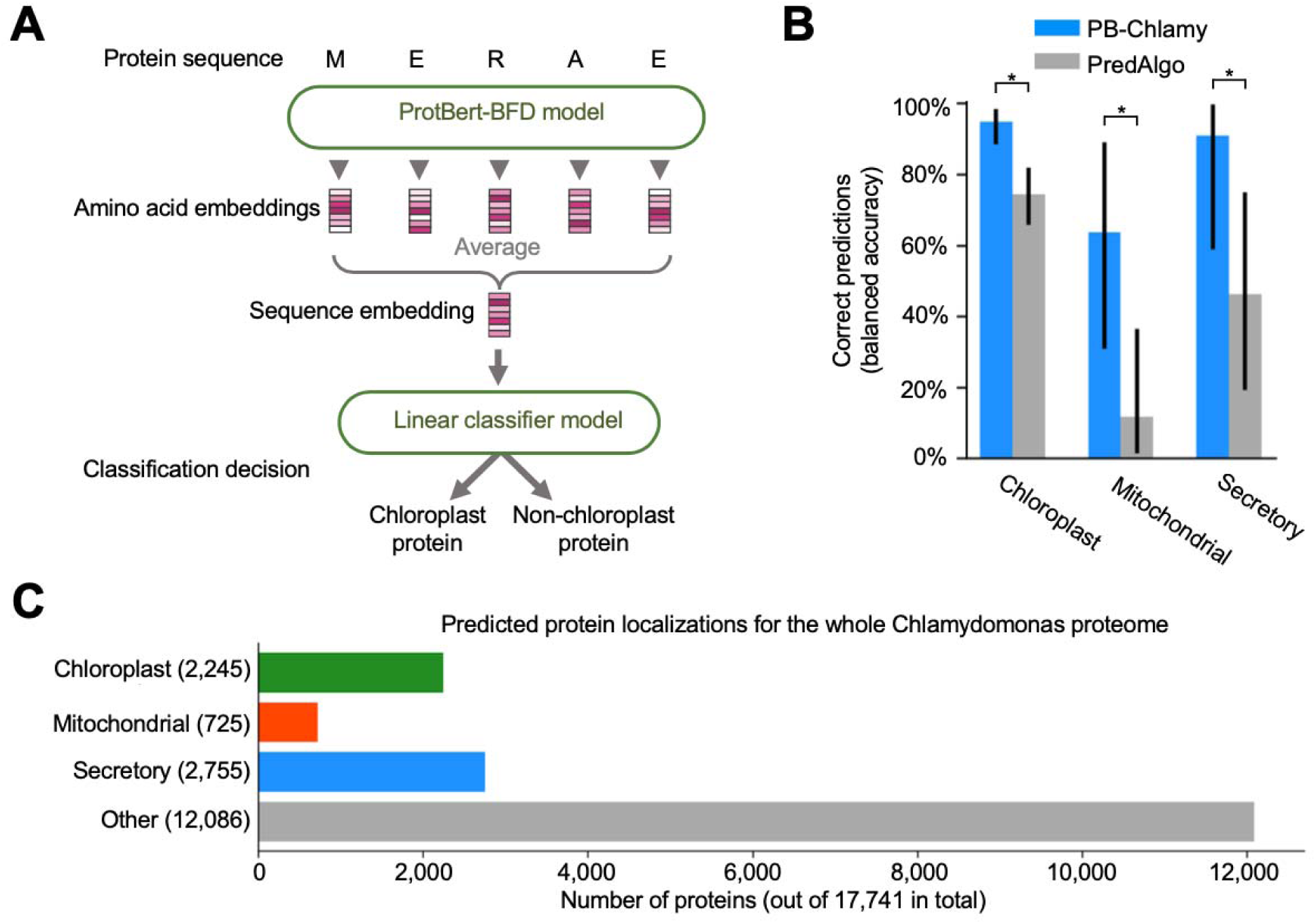
We predicted localizations of all proteins in *Chlamydomonas reinhardtii*. (A) The PB-Chlamy set of predictors was created based on a pre-trained ProtBertBFD instance with second-stage supervised training using our protein localization dataset. (B) PB-Chlamy displays improved accuracy compared to PredAlgo, the previous state-of-the-art localization predictor for Chlamydomonas, on a test set randomly chosen from our localized proteins and not used for training. The measure displayed is balanced accuracy, an average of sensitivity (the % of real positives correctly classified) and specificity (the % of real negatives correctly classified). Each separate localization predictor had its own test set, with sizes as follows: 111 chloroplast and 97 non-chloroplast proteins, 12 mitochondrial and 194 non-mitochondrial, 18 secretory and 149 non-secretory. The asterisk * indicates p<0.05. (C) PB-Chlamy was used to predict localizations for the whole Chlamydomonas proteome (v5.6, primary transcripts only).

We evaluated the performance of PB-Chlamy in comparison to PredAlgo. To ensure that neither of the predictors being compared had been trained on any of the proteins in the test sets, we used testing datasets with proteins used to train PredAlgo excluded. PB-Chlamy reliably performs better than PredAlgo on our test sets for proteins localized to the chloroplast, mitochondria, and secretory pathway (Figure 7B and S7).

We proceeded to use PB-Chlamy to predict protein localizations for the entire Chlamydomonas proteome (Figure 7C; Table S7), finding 2,245 putative chloroplast proteins, 725 putative mitochondrial proteins, and 2,755 putative secretory proteins. These numbers include 70 proteins with predicted dual localizations, mostly chloroplast+mitochondria (Table S7). Notably, we predict only two-thirds as many chloroplast proteins and one-quarter as many mitochondrial proteins as PredAlgo (which predicts 3,375 chloroplast and 2,843 mitochondrial proteins), providing a sharper view of the predicted proteome of these organelles.

## DISCUSSION

### Insights into the Functions of Poorly-Characterized Proteins

Our localization information is the first compendium of this kind in any photosynthetic organism. The data are particularly useful for advancing the understanding of the molecular functions of poorly-characterized proteins. The molecular functions of 702 (68%) of our localized proteins are unknown and 459 (45%) of the localized proteins were previously unnamed (Figure S1F). We provided examples of how our localization data give insights into the functions of such poorly-characterized proteins, like the predicted DNA-binding protein SND1B, which we localized to chloroplast nucleoids. We also illustrated of how the function of poorly-characterized proteins can be elucidated by immunoprecipitation and mass spectrometry of our tagged strains, as in the case of the poorly-characterized proteins PGC1 and PGC2 where our data allowed us to assign them to plastoglobules. For proteins where we could not obtain experimental localization data, our new PB-Chlamy classifier accurately predicts their localization. Together, our localization data, protein-protein interactions, and computational predictions greatly narrow down the possible functions of poorly-characterized proteins and facilitate generation of specific hypotheses for their further characterization, accelerating the elucidation of chloroplast organization and function.

The images and protein-protein interactions from this study can be browsed and searched at https://www.chlamylibrary.org/. This site also provides links for ordering the corresponding strains and plasmids from the Chlamydomonas Resource Center.

### Novel Organizational Features of the Chloroplast

Our systematic survey of protein localizations revealed extensive spatial organization of the chloroplast. This organization includes 11 novel punctate structures, which appear to be metabolic hubs that enhance or regulate specific reactions such as the commitment step of L-serine biosynthesis or the final step of chlorophyll biosynthesis. We observed the enrichment of Calvin-Benson cycle enzymes in the proximity of the pyrenoid, which may enhance the cycle’s overall activity by localizing each enzyme to the site where its substrate is produced. We observed extensive spatial organization of the thylakoid membrane and chloroplast envelope, including distinct chloroplast envelope regions that appear to be specialized for interactions with cytosolic ribosomes, and other regions that appear to be specialized for interactions with chloroplast ribosomes. These discoveries open doors to characterizing the functions of these organizational features and of the molecular bases that underlie their organization.

### Insights Into Dual Targeting

Our study provides the largest-scale survey to date of proteins with multiple compartment localization in any photosynthetic eukaryote, making it a resource for studying the biological functions of this phenomenon, the mechanisms that underlie it, and its evolution. Whereas the vast majority of previously-known dual-localized plant proteins were dual-localized to the chloroplast and mitochondria (Carrie and Whelan, 2013), only two of our proteins showed this localization and the vast majority of our dual-localized chloroplast proteins had a second localization in a compartment other than mitochondria (Figure 6B and 6C; Table S2), making our dataset a complementary resource. We observed dual localization to the chloroplast and a broad range of other organelles, including previously un-described dual localizations to crescent-shaped structures that we propose based on corroborating data to correspond to acidocalcisomes. These observations suggest widespread sharing of functions between the chloroplast and cytosolic organelles, and/or extensive cytosolic degradation of chloroplast proteins. Our identification of these dual-localized proteins provides a starting point for future characterization of the functional and regulatory interactions between the corresponding organelles.

### Evolution of Chloroplast-Associated Protein Localizations and Interactions

Our study provides insights into both genes specific to the Chlorophyte green algal lineage and genes conserved across the eukaryotic supergroup *Archaeplastidia*: 933 of the Chlamydomonas proteins we localized are conserved in the green alga *Volvox carteri*, 696 are conserved in the green alga *Coccomyxa subellipsoidea*, and 618 are conserved in the land plant *Arabidopsis thaliana* (Figure S1G; Table S2). We observed similarities and differences in protein localizations between Chlamydomonas and land plants for proteins that show single and multiple localizations (Table S2). For example, the Chlamydomonas protein Cre12.g494250 localized to the chloroplast, cytosol, nucleus, and mitochondrion; and its Arabidopsis homolog AT4G16060 showed a similar multiple localization to the chloroplast, cytosol, and mitochondrion when expressed in tobacco leaves (Figure 6T, S6W, and S6X). These observations suggest that our dataset provides a rich resource that will allow the study of the evolutionary principles of protein localization and of dual targeting.

### Perspective

Climate change and the rising global population drive a pressing need to understand the basic biology of photosynthetic organisms and to advance our ability to engineer them. Our study lays the groundwork for understanding remaining mysteries of the chloroplast, the organelle at the heart of photosynthetic organisms. Importantly, the systematic and comprehensive nature of the present study allowed us to reach deep into the unknown, revealing organizational features that would not have been readily accessible with traditional hypothesis-driven research. We hope that further characterization by the research community of these data and strains will help advance the understanding of the evolution of protein localization, the roles of poorly-characterized chloroplast proteins, and the spatial organization of chloroplast functions.

## SUPPLEMENTAL INFORMATION

Supplemental Information includes 7 figures, 8 tables, and 1 movie.

## AUTHOR CONTRIBUTIONS

L.W. and M.C.J. conceived the project. L.W., K.A.V.B., Y.X., S.G., and H.H. performed the gene cloning, generating fluorescently tagged strains, and confocal microscopy. M.W. prepared materials for this large-scale work. L.W. and E.R.S. performed immunoprecipitation and prepared the samples for analysis by mass spectrometry. S.K. and H.H.S conducted the mass spectrometry. W.P. conducted bioinformatic prediction of protein localizations. C.D.M., V.C., and V.T. conducted the prediction of protein structure. L.W. performed indirect immunofluorescence assay. L.H., D.J.S., C.B., and J.H. performed the protein localization in Tobacco. M.H.C., W.P., L.W., and M.C.J. built the website to share the data and materials. All authors analyzed the data. L.W., A.T.W., and M.C.J. wrote the manuscript with input from all authors.

## STAR*METHODS

### KEY RESOURCES TABLE

| REAGENT or RESOURCE | SOURCE | IDENTIFIER |
| --- | --- | --- |
| **Antibodies** |  |  |
| Goat anti-Rabbit IgG(H+L) Highly Cross-Adsorbed Secondary Antibody, Alexa Fluor 488 | Invitrogen | Cat# A11034 |
| Rabbit polyclonal anti-Cre01.g028150 | This paper | N/A |
| Rabbit polyclonal anti-Cre01.g013150 | This paper | N/A |
| Rabbit polyclonal anti-Cre12.g519350 | This paper | N/A |
| Rabbit polyclonal anti-Cre03.g172550 | This paper | N/A |
| Rabbit polyclonal anti-Cre01.g019250 | This paper | N/A |
| **Chemicals, Peptides, and Recombinant proteins** |  |  |
| Digitonin, Water soluble cOmplete, EDTA-free Protease inhibitor | Research Products International Roche | Cat# 11024-24-1<br>Cat# 5056489001 |
| Anti-FLAG M2 Magnetic Beads | Sigma-Aldrich | Cat# M8823 |
| 3xFLAG peptide | Sigma-Aldrich | Cat# F4799 |
| 4xLaemmli sample buffer | Bio-Rad | Cat# 1610747 |
| Dimethyl sulfoxide | Sigma-Aldrich | Cat# 67-68-5 |
| Betaine | Sigma-Aldrich | Cat# 107-43-7 |
| Formaldehyde solution 4 %, pH6.9 | Sigma-Aldrich | Cat# 1004960700 |
| Bovine Serum Albumin | Sigma-Aldrich | Cat# A3059 |
| VECTASHIELD Mounting Medium | Vector Laboratories | Cat# H-1300 |
| UltraPure Low-Melting Point Agarose | Invitrogen | Cat# 16500100 |
| 4-15 % Criterion TGX Precast Midi Protein Gel | Bio-Rad | Cat# 5671084 |
| MAX efficiency transformation Reagent for Algae | Invitrogen | Cat# A24229 |
| DL-Dithiothreitol | Sigma-Aldrich | Cat# D0632 |
| Trypsin Gold, Mass Spectrometry Grade | Promega | Cat# V5280 |
| **Critical Commercial Assays** |  |  |
| Phusion High-Fidelity DNA polymerase | New England BioLabs | Cat# M0530L |
| MinElute Gel Extraction Kit | QIAGEN | Cat# 28606 |
| Gibson Assembly Master Mix | New England BioLabs | Cat# E2611L |
| Invitrogen Gateway Clonase II | Invitrogen | Cat#11791020 |
| QIAprep Spin Miniprep Kit | QIAGEN | Cat# 27106 |
| **Experimental Models: Organisms/Strains** |  |  |
| *C.reinhardtii*: wild-type CC-4453 | Chlamydomonas Resource Center | CC-4533 cw15 |
| *E. coli* Stellar Competent Cells | Takara | Cat# 636763 |
| Chlamydomonas strains expressing tagged proteins listed in Table S6 | This paper, Chlamydomonas Resource Center | https://www.chlamycollection.org/ |
| **Oligonucleotides and Recombinant DNA** |  |  |
| pLM005 | (Mackinder et al., 2016); GenBank | KX077945.1 |
| pLM006 | (Mackinder et al., 2016); GenBank | KX077949.1 |
| pENTR223 | ABRC | https://www.arabidopsis.org/ |
| pEarleyGate 103 | ABRC | https://www.arabidopsis.org/ |
| Mitochondria marker (CD3-991) | ABRC | https://www.arabidopsis.org/ |
| Plasmid constructs generated and listed in Table S1 | This paper, Chlamydomonas Resource | https://www.chlamycollection.org/ |
| Software and Algorithms |  |  |
| Fiji | (Schindelin et al., 2012) | https://imagej.net/software/fiji/downloads |
| Thermo Proteome Discoverer 2.5 | Thermo Scientific | Cat# OPTON-30945 |
| *Continued* |  |  |
| REAGENT or RESOURCE | SOURCE | IDENTIFIER |
| Scaffold 5 | Proteome Software | https://www.proteomesoftware.com/products/scaffold-5 |
| Others |  |  |
| Electroporation Cuvette, 2mm gap | Bulldog Bio. | Cat# 12358-346 |
| Ibidi USA μ-Slide 8 well, Glass bottom | Ibidi | Cat# NC0704855 |
| Poly-L-lysine coated glass slides | Sigma-Aldrich | Cat# P0425 |
| Kontes Duall #22 homogenizer | Kimble | Cat# KT885450-0022 |
| Avanti J-26X with 8.1000 rotor | Beckman Coulter | N/A |
| Cryomill | Retsch | Part NO. 20.749.0001 |
| Electroporator | NEPA GENE | NEPA21 type II |
| SP5 Confocal Microscope | Leica | TCS SP5 |
| Singer Rotor HAD | Singer Instruments | Cat# ROT-001 |
| Typhoon FLA9500 fluorescence scanner | GE Healthcare | N/A |
| Colony picker | Norgren Systems | Part NO.12461 |
| Nikon Confocal laser scanning microscope | Nikon | A1Rsi |
| Nikon Confocal microscope | Nikon | A1R-STED |

### CONTACT FOR REAGENT AND RESOURCE SHARING

Further information and requests for resources and reagents should be directed to and will be fulfilled by the Lead Contact, Martin C. Jonikas

### EXPERIMENTAL MODEL AND SUBJECT DETAILS

#### Strains and culture conditions

The *Chlamydomonas reinhardtii* strain CC-4533 (cMJ030) was used for wild-type (hereafter WT) in all experiments. All strains were maintained on Tris-acetate-phosphate (TAP) solid medium with 1.5 % agar at 22 °C under dim light (<10 μmol photons m^-2^s^-1^). All media used revised trace element solution (Kropat et al., 2011).

### METHOD DETAILS

#### Target genes selection

Target genes were selected from seven sources (Figure S1A), including 1,093 genes encoding proteins identified in Arabidopsis chloroplast proteomics (Ferro et al., 2010), 644 genes encoding proteins identified in Chlamydomonas chloroplast proteomics (Terashima et al., 2010), 154 genes encoding proteins identified in Chlamydomonas pyrenoid proteomics (Zhan et al., 2018), 3,317 genes encoding PredAlgo predicted chloroplast proteins and 858 genes encoding proteins with low PredAlgo score in non-chloroplast organelles (Tardif et al., 2012), 510 genes encoding GreenCut2 proteins (Karpowicz et al., 2011), and 303 genes encoding candidate proteins required for photosynthesis suggested in mutant screening (Li et al., 2019). In addition, we also selected 777 genes because of their potential association with chloroplast function suggested either in their Phytozome annotation (https://phytozome-next.jgi.doe.gov/) or in related reports, such as the TEF proteins present in thylakoid enriched fraction (Allmer et al., 2006) and FTT proteins interacting with well-known chloroplast proteins (Mackinder et al., 2017). To avoid duplicating effort, we removed the overlapping genes across the seven sources above and genes encoding proteins which had been localized in Mackinder et al., 2017. Altogether, we obtained 5,874 target genes.

#### Plasmid Construction and Cloning

We designed our primers according to gene sequences present in the v5.5 *Chlamydomonas reinhardtii* genome. Cross et al. (Cross, 2016) identified upstream ATGs in many of these gene sequences, and supplementary data in Mackinder et al. 2017 indicate that for genes that include such upstream ATGs, using the original ATG leads to lower localization success rates, suggesting that the Cross et al. 2016 upstream ATGs more frequently correspond to the native translation start site. Therefore, wherever an upstream ATG had been identified by Cross et al., we used this ATG instead of the one annotated in the genome, leading to our usage of a corrected upstream ATG in 1,213 of our target genes (Table S1).

The cloning pipeline was based on that used in Mackinder et al. 2017, with some modifications. The open reading fames were amplified from Chlamydomonas WT genomic DNA by PCR using Phusion High-Fidelity DNA polymerase (New England BioLabs) with additives of 6 % DMSO (v/v) (Sigma-Aldrich) and 1 M Betaine (Sigma-Aldrich). The PCR products were gel purified using MinElute Gel Extraction Kit (QIAGEN) and then cloned in-frame with a C-terminal Venus-3×FLAG in pLM005 by Gibson assembly (New England BioLabs). Primers were designed to amplify the open reading frame until but excluding the stop codon, and with adaptors to allow efficient assembly into *Hpal*-cut pLM005. Considering the PCR limitations to amplification of large genes, we mainly focused on genes smaller than 8 kb in this study. For genes larger than 6 kb, we split them into multiple fragments (< 3 kb) for PCR amplification and then reassembled the fragments together during the final Gibson assembly step. The fragment size was verified by restriction enzyme digestion. A pilot study showed that 334/334 (100 %) of genes had correct junctions as verified by Sanger sequencing.

Cloning of Chlamydomonas genes is known to be challenging due to high GC content, repetitive sequences, and gene length (Mackinder et al., 2017). In total, we successfully cloned 3,116 genes (53 %) (Figure S1A), a similar fraction to the 48 % in Mackinder et al. 2017. Interestingly, the cloning success of genes smaller than 500 bp is 66.2 %, which is lower than 86.5 % of genes with size between 1,000∼2,000 bp (Figure S1H).

#### Chlamydomonas transformation

Constructs were linearized by *EcoRV*, *DraI*, *AfIII*, or *Bsal* prior to the electroporation into WT Chlamydomonas strain CC-4533. WT cells were pre-cultured in TAP liquid medium at 22 °C under light with a photon flux density of 150 μmol photons m^−2^ s^−1^ until the cell density reached to ∼2 ×10^6^ cells mL-1.For each transformation, 150 ng of cut plasmid was mixed with 60 μL of 2×10^8^ cells mL^-1^ suspended in MAX Efficiency Transformation reagent (Invitrogen) in an ice-cold 0.2 cm gap electroporation cuvette (Bulldog Bio.) and transformed into WT strains by electroporation using a NEPA21 electroporator (NEPA GENE) (Yamano et al., 2013). The settings were: Poring Pulse: 250.0 Volts, 8.0 ms pulse length, 50.0 ms pulse interval, 2 pulses, 10 % decay rate, + polarity; Transfer Pulse: 20.0 Volts, 50.0 ms pulse length), 50.0 ms pulse interval), 10 pulses, 40 % decay rate), +/- polarity. For recovery, cells were transferred to 10 mL TAP liquid medium plus 40 mM sucrose and incubated with gentle shaking under dim light (<10 μmol photons m^-2^s^-1^) overnight. The transformants were plated on TAP agar medium supplied with 20 μg mL^-1^ paromomycin under light (150 μmol photons m^−2^ s^−1^.). After 7 days incubation under dim light (<10 μmol photons m^−2^ s^−1^), 48 transformants from each plate were arrayed on a new rectangular TAP agar PlusPlate (Singer Instruments) using a colony Picker (Norgren Systems). The transformants were replicated manually onto a fresh TAP agar PlusPlate using a 96-Long pin pad (Singer Instruments). The TAP plates with arrayed transformants were screened for fluorescence using a Typhoon FLA9500 fluorescence scanner (GE Healthcare) with the following settings: Venus, 532 nm excitation with 555/20 nm emission. The colonies with positive fluorescence signals were isolated and maintained in 96 arrays using a Singer Rotor propagation robot (Singer Instruments).

Transformation of constructs and localization of proteins in Chlamydomonas are known to be inefficient (Mackinder et al., 2017), possibly due to several mechanisms that fight foreign DNA (Neupert et al., 2009; Zhang et al., 2014). Our transformation and localization success rate (34 %) was lower than that in Mackinder et al. 2017 (49 %), possibly because the genes targeted in the present study were overall expressed at lower levels. To generate dual-tag lines, pLM006 harboring an mCherry-6×HIS tag was used as the backbone, and TAP agar medium supplied with 20 μg mL^-1^ hygromycin was used for selection.

#### Confocal Microscopy

For confocal imaging, colonies were transferred to a 96-well microtiter plate with 100 μL TP liquid medium and 5 μg mL^-1^ antibiotics in each well and then pre-cultured in air under 150 μmol photons m^−2^ s^−1^ on an orbital shaker with gentle agitation of 600 RPM. After ∼16 hr of growth, 10 μL cells were transferred onto an μ-Slide 8-well glass-bottom plate (Ibidi) and 200 μL of 1 % TP low-melting-point agarose at ∼35 °C was overlaid to restrict cell movement. All imaging except for Fluorescence Recovery After Photobleaching (FRAP) assays was conducted using a Leica SP5 confocal microscope with the following settings: Venus, 514 nm excitation with 530/10 nm emission; mCherry, 561 nm excitation with 610/30 nm emission; and chlorophyll, 514 nm excitation with 685/40 nm emission. All confocal microscopy images were analyzed using Fiji (Schindelin et al., 2012). For each strain, a confocal section through a cell showing the predominant localization pattern was captured and analyzed. To minimize the bias in determining the localization patterns, each localization image was independently analyzed by two researchers. Localization patterns for 31 proteins where there was clear disagreement or insufficient signal were categorized as Ambiguous.

FRAP assays were performed using a Nikon A1R-STED confocal microscope with the following setting: Venus 514 nm excitation with 530/10 nm emission; and chlorophyll, 514 excitation with 685/40 nm emission. One baseline image was acquired before FRAP was performed. The selected puncta were bleached by a high-intensity laser beam (514 nm wavelength). The recovery of fluorescence at the bleached puncta was imaged every 30 s for 10 min.

#### Indirect Immunofluorescence Assay

Indirect immunofluorescence was performed as described previously (Wang et al., 2016). Briefly, Cells were harvested by centrifugation and rinsed with PBS buffer twice. Then 100 μL of cells was spotted onto Poly-L-lysine coated glass slides (Sigma-Aldrich). Cells were fixed with 4 % (w/v) formaldehyde (Sigma-Aldrich) in PBS for 20 min and then incubated with 100 % ice-cold methanol for 20 min to remove chlorophyll. Purified antibodies (Yenzyme) against Cre01.g028150, Cre01.g013150, Cre12.g519350, Cre03.g172550, and Cre01.g019250, were used at a dilution of 1:200. The purified antibodies were generated using the following peptides: C-Ahx-PDQPPRILTTRRE-amide (Cre01.g028150), C-Ahx-TWDVKAPINKHYNFH-cooh (Cre01.g013150), C-Ahx-YLPNTGNMLMQVNPNQ-cooh (Cre12.g519350), C-Ahx-RGQVKNTQQYRMR-cooh (Cre03.g172550), and C-Ahx-KGVDATKYSHSTIVQT-amide (Cre01.g019250). After washing the slides 4 times, each with 50 mL PBS-T (supplied with 0.1% Tween 20 (v/v)) in Coplin jar, Alexa Fluor 488 goat anti-rabbit IgG (H+L) Cross-Adsorbed Secondary Antibody (Invitrogen) was used at a dilution of 1:500. Then, washing the slides 4 times, each with 50 mL PBS-T. Fluorescence and bright-field images were acquired using a confocal microscope (Leica, SP5).

#### Protein Structure Modeling

Alphafold2.1 (Jumper et al., 2021) was used to screen for candidate multimeric structures of proteins identified in pull-down experiments of plastoglobule complex, using one NVIDIA A100 GPU. Pairs of protein sequences were concatenated in silico with a 50-U linker sequence (Moriwaki, 2021), and structures were predicted with Alphafold-monomer model. Proteins found to dimerize were again compared to other proteins in the same pull-down experiment. ChimeraX (Pettersen et al., 2021) was used to visualize structures. The Alphafold2 predicted complex of Cre02.g143667 and Cre06.g286300 was deposited in Modelarchive (https://www.modelarchive.org/).

#### Protein localization prediction

For each subcellular localization, we trained a protein language model to predict protein localization from protein sequence. Protein language models are first trained on large numbers of sequences, and then these pretrained models can be retrained for a specific prediction task, in this case subcellular location prediction (Figure 7A). For our pretrained model, we used ProtBertBFD, a protein language model pre-trained on billions of protein sequences (Elnaggar et al., 2021; https://huggingface.co/Rostlab/prot_bert_bfd). Given a protein sequence, ProtBertBFD outputs numeric vectors, or embeddings, that capture features of each amino acid in that sequence. These amino acid embeddings contain information on biochemical and structural properties (Elnaggar et al., 2021). To represent a protein sequence, we take an average of the embeddings of all of its amino acids, and use this representation as an input to a linear classifier to distinguish if a protein is localized to a particular cellular compartment or not. Specifically, we use the model architecture BertForSequenceClassification from the huggingface python package (Wolf et al., 2020). Our script for running training and evaluation is https://github.com/clairemcwhite/transformer_infrastructure/hf_classification.py. For each compartment (chloroplast, mitochondrial and secretory) we used proteins found to localize to the compartment as positive cases, and proteins not found to localize to the compartment as negative cases. We used a random 60% of these positive and negative cases to train the model, 20% for performance validation during training, and 20% as a fully withheld test set to evaluate model performance on unseen examples. These sets are listed in Table S7.

The raw score distributions, PR and ROC curves and summary measures compared to PredAlgo are shown in Figure S11; for the purpose of comparisons with PredAlgo, we used the testing sets minus any proteins that were included in PredAlgo training data. We then used the trained models to predict protein localizations for the entire Chlamydomonas proteome (Table S7).

We downloaded Chlamydomonas protein sequences from Phytozome (https://phytozome-next.jgi.doe.gov/info/Creinhardtii_v5_6, genome version 5.6); we only used primary transcripts for training and for localization prediction. We adjusted the protein sequences to use the new start codons described by Frederick Cross (Cross, 2016) - they are included in Table S7.

The training command and environment setup for the chloroplast were as follows, with analogous commands for the other localizations:

module load cudatoolkit

bash make_hf-transformers_conda_env.sh

conda activate hf-transformers

python hf_classification.py -m prot_bert_bfd/ -tr chloro_train.csv -v chloro_val.csv –te

chloro_test.csv -o results_chloro -maxl 1150 -n chloro -e 10 -tbsize 1 -vbsize 1 -s 3

The input files containing the training/validation/test sets were plaintext, formatted as follows: Entry_name,sequence,label

Cre03.g155400,M H K T P C L H G G S S L S A G R A P L A R L C C A S Q R V R G P A P A Q A F W K Q S G A S A G K S G K A R P G A K A Q Q P K Q K A G G G K Q G G G G G G G L M D S E V P V Y A E A F D I N K C V D L Y L R F F K W V S S P V T G G S G K

K,Chloroplast

The trained model files are available (https://huggingface.co/wpatena/PB-Chlamy/tree/main).

#### Affinity Purification and Mass Spectrometry

Each affinity purification-mass spectrometry (AP-MS) experiment was performed twice from independently grown samples of the same strain. Cells expressing Venus-3×FLAG-tagged proteins were pre-cultured in 50 mL TAP medium with 5 μg mL^-1^ paromomycin until the cell density reach to ∼2-4 ×10^6^ cells mL^-1^. Then, cells were harvested by centrifugation at 1,000 g for 5 min and the pellets were suspended in 1,000 mL TP liquid medium. Cells were grown with air bubbling and constant stirring of 210 RPM under 150 μmol photons m^−2^ s^−1^ light until the cell density reached ∼2-4 ×10^6^ cells mL^-1^. Cells were collected by centrifugation at 3,000 g for 4 min in an Avanti J-26X centrifuge with an 8.1000 rotor (Beckman) at 4 °C. The pellets were washed in 35 mL ice-cold washing buffer (25 mM HEPES, 25 mM KOAc, 1 mM Mg(OAc)_2_, 0.5 mM CaCl_2_, 100 mM Sorbitol, 1mM NaF, 0.3 mM Na_3_VO_4_, and cOmplete EDTA-free protease inhibitor (1 tablet/500 mL)) and then resuspended in a 1:1 (v/w) ratio of ice-cold 2×IP buffer (50 mM HEPES, 50 mM KOAc, 2 mM Mg(OAc)_2_, 1 mM Cacl_2_, 200 mM Sorbitol, 1mM NaF, 0.3 mM Na_3_VO_4_, and cOmplete EDTA-free protease inhibitor (1 tablet/50 mL)). 3 mL cell slurry was immediately added to liquid nitrogen to form small popcorn pellets which were stored at -80 °C until needed. Cells were lysed by cryogenic grinding using a Cryomill (Retsch) at frequency of 25 oscillations per second for 20 min. The ground powder was defrosted on ice for 45 min and dounced 25 times on ice with a Kontes Duall #22 homogenizer (Kimble). 1mL homogenized cells of each sample was used for the following processes. Membrane proteins were solubilized by incrementally adding an equal volume of ice-cold 1×IP buffer plus 2 % digitonin (RPI) followed by an incubation of 45 min with nutation at 4 °C. The cell debris were removed by spinning at 12,700 g for 30 min at 4°C. The supernatant was then mixed with 50 μL anti 3×FLAG magnetic beads (Sigma) which had been previously washed sequentially with 1×IP buffer 3 times and 1×IP buffer plus 0.1 % digitonin 2 times. The mixture was incubated with nutation at 4 °C for 1.5 hr, followed by the removal of supernatant. The beads were washed 4 times with 1×IP buffer plus 0.1 % digitonin followed by a 30 min competitive elution with 45 μL of 1×IP buffer plus 0.25 % digitonin and 2 μg/μL 3×FLAG peptide (Sigma-Aldrich). After elution, 30 μL protein samples were mixed with 9.75 μL 4×SDS-PAGE buffer (Bio-Rad) containing 100 mM DTT (Sigma-Aldrich) followed by denaturation by heating at 70 °C for 10 min. Then, 30 μL denatured protein sample was loaded into a well of a 4-15 % Criterion TGX Precast Midi Protein Gel (BioRad) for electrophoresis at 50 V for 40 min until the protein front moved ∼2.5 cm. ∼2.0 cm gel slice containing target proteins with molecular weight >= 10 kDa (to exclude the 3xFLAG peptide) were excised and stored at 4 °C until processing for in-gel digestion. To decrease cross-contamination from samples in neighboring wells, samples were loaded in every other well. To further avoid carry-over contamination of mass spectrometry and contamination from sequential samples, we performed two biological repeats of AP-MS and changed the order of samples in the two biological repeats.

In-gel digestion of protein bands using trypsin was performed as previously (Shevchenko et al., 2006). Trypsin digested samples were dried completely in a SpeedVac and resuspended with 20 μL of 0.1 % formic acid pH 3 in water. 2 μL (∼ 360 ng) was injected per run using an Easy-nLC 1,200 UPLC system. Samples were loaded directly onto a 15 cm long, 75 μm inner diameter nanocapillary column packed with 1.9 μm C18-AQ resin (Dr. Maisch, Germany) mated to a metal emitter in-line with an Orbitrap Fusion Lumos (Thermo Scientific, USA). The column temperature was set at 45 °C and a half-hour gradient method with 300 nL per minute flow was used. The mass spectrometer was operated in data dependent mode with a 120,000 resolution MS1 scan (positive mode, profile data type, AGC gain of 4e5, maximum injection time of 54 s and mass range of 375-1,500 m/z) in the Orbitrap followed by HCD fragmentation in ion trap with 35 % collision energy. A dynamic exclusion list was invoked to exclude previously sequenced peptides for 60s and a maximum cycle time of 3 s was used. Peptides were isolated for fragmentation using the quadrupole (1.2 m/z isolation window). The ion trap was operated in Rapid mode.

#### Transient expression of Arabidopsis gene in Tobacco leaf

We first cloned the full-length cDNA of AT4g16060 into pENTR223 (ABRC). Then the AT4G16060 was cloned in-frame with a C-terminal GFP in pEarleyGate 103 by LR recombination reaction (Invitrogen Gateway Clonase II). The construct of AT4G16060-GFP and mitochondria mCherry marker CD3-991 (ABRC) were then separately transformed into Agrobacterium tumefaciens by heat shock. A mixture (OD600 = 0.125) of the Agrobacterium tumefaciens harboring At4G16060-GFP, the Agrobacterium tumefaciens carrying CD3-991, and the p19 protein of tomato bushy stunt virus (TBSV) in a 2:1:2 ratio co-infiltrated into 4-weeks old tobacco plants (*Nicotiana tabacum*) (Sainsbury et al., 2009). Three days after infiltration, the abaxial epidermis of the leaves were imaged using Nikon A1Rsi confocal laser scanning microscopy (CLSM) with Nikon 60x Apo (NA1.40) objective. The imaging was performed with the following setting: GFP, 488 nm excitation with 525/25 nm emission; mCherry, 561 nm excitation with 595/25 nm emission; and Chlorophyll, 647 nm excitation with 699/37 nm emission. Image acquisition and analysis were performed using the Nikon NIS Elements software (version 5.21.03).

### QUANTIFICATION AND STATISTICAL ANALYSIS

#### Peptide identification

Raw files were searched using MSAmanda 2.0 (Dorfer et al., 2021) and Sequest HT algorithms (Eng et al., 1994) within the Proteome Discoverer 2.5.0 suite (Thermo Scientific, USA). 10 ppm MS1 and 0.4 Da MS2 mass tolerances were specified. Carbamidomethylation of cysteine was used as fixed modification, oxidation of methionine, deamidation of asparagine and glutamine were specified as dynamic modifications. Pyro glutamate conversion from glutamic acid and glutamine are set as dynamic modifications at peptide N-terminus. Acetylation was specified as dynamic modification at protein N-terminus. Trypsin digestion was selected with a maximum of 2 missed cleavages allowed. Files were searched against the UP000006906 Chlamydomonas database downloaded from Uniprot.org (https://www.uniprot.org/).

Scaffold (version Scaffold 5.1.0, Proteome Software Inc., Portland, OR) was used to validate MS/MS based peptide and protein identifications. Peptide identifications were accepted if they could be established at greater than 95.0 % probability by the Scaffold Local FDR algorithm. Protein identifications were accepted if they could be established at greater than 99.9 % probability and contained at least 2 identified peptides. Protein probabilities were assigned by the Protein Prophet algorithm (Nesvizhskii et al., 2003).

#### Calculating WD-scores

The WD-scores of MS data were calculated using the ComPASS method, which analyzes spectral counts based on the specificity of the prey, spectral count number and reproducibility (Sowa et al., 2009). Instead of using the spectral counts from two technical repeats, we used the spectral counts from two biological replicas with different neighbors for each sample. First, we generate a Stats table containing all the bait proteins and interactors as below,

| | Bait 1 | Bait2 | Bait3 | Bait4 | Bait $k$ | |
| --- | --- | --- | --- | --- | --- | --- |
| Interactor 1 | $X_{1,1}$ | $X_{2,1}$ | $X_{3,1}$ | $X_{4,1}$ | $X_{k,1}$ | $\bar{X}_1$ |
| Interactor 2 | $X_{1,2}$ | $X_{2,2}$ | $X_{3,2}$ | $X_{4,2}$ | $X_{k,2}$ | $\bar{X}_2$ |
| Interactor 3 | $X_{1,3}$ | $X_{2,3}$ | $X_{3,3}$ | $X_{4,3}$ | $X_{k,3}$ | |
| Interactor 4 | $X_{1,4}$ | $X_{2,4}$ | $X_{3,4}$ | $X_{4,4}$ | $X_{k,4}$ | |
| Interactor $m$ | $X_{1,m}$ | $X_{2,m}$ | $X_{3,m}$ | $X_{4,m}$ | $X_{k,m}$ | $\bar{X}_m$ |

X*_i_*_,*j*_ is the average spectral counts from two biological replicas for interactor *j* from bait *i*. *m* is the total number of unique prey proteins identified (8,067).

*k* is the total number of unique bait (41).

We calculated the WD-scores using the equations ((Sowa et al., 2009)) below,

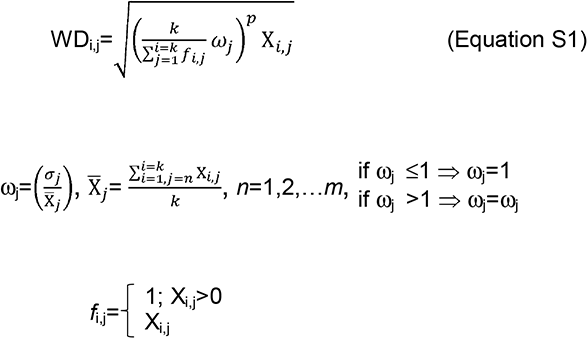

*p* is the number of replicates runs in which the interactor is present *f_i,j_*.

The minimum WD score values for high-confidence interactions will be different for each study because the WD score depends on the specific proteins and methods used in each study. In Mackinder et al., 2017, we set the high-confidence WD cut-off based on WD-scores of prey proteins that localized to a different compartment than the bait. Because all the baits in the present study localized to the same compartment (the chloroplast) we could not use the same approach to set the WD cut-off. We therefore set our WD cutoff at 3.7 % of all interactions based on the corresponding value in Mackinder et al., 2017 of 3.78 %. This rationale led to a WD cut-off for the present study of 16.567, with 297 of the 8,067 interactions above this threshold (Table S5). We defined high-confidence protein-protein interactions as those having a WD score above the cutoff of 16.567 and where the prey was detected in both biological repeats, which resulted in ∼187 high-confidence protein-protein interactions (Table S5).

#### Data visualization

The calculation of WD-score and assembly of bait-prey matrix were performed in Microsoft Exel. The alignment of amino acid sequence was conducted using Clustal Omega with default settings (https://www.ebi.ac.uk/Tools/msa/clustalo/). The protein structures were visualized using ChimeraX (https://www.cgl.ucsf.edu/chimerax/).

#### Transmembrane prediction and Protein homology prediction

Protein transmembrane domains were predicted using TMHMM2.0 (https://services.healthtech.dtu.dk/service.php?TMHMM-2.0). Protein homologies were predicted using Phyre2 (http://www.sbg.bio.ic.ac.uk/phyre2/html/page.cgi?id=index)

#### Statistical tests

Statistical tests comparing PredAlgo and PB-Chlamy were performed in Python using scipy.stats and rpy2. All other statistical tests were performed in Microsoft Excel.

### ADDITIONAL RESOURCES

Protein localization images are available at https://www.chlamylibrary.org/.

Fluorescently tagged strains and plasmid constructs are available at https://www.chlamycollection.org/.

**Table S1. List of the 5,784 target genes that we attempted to clone and localize in this study**. Related to Figure 1 and Star Methods.

**Table S2. Localizations of 1,032 proteins localized in this study.** Related to Figure 1.

**Table S3. Number of proteins exhibiting each of the 141 distinct localization patterns**. Related to Figure 1.

**Table S4. Our localizations match previously-published localizations for 27 of the 28 previously-localized Chlamydomonas proteins represented in our study.** Related to Figure 1.

**Table S5. Whole list of protein-protein interactions identified in this study.** Related to Figure 1-6.

**Table S6. Links for viewing the data on Chlamylibrary.org and information for ordering plasmids and strains from the Chlamydomonas Resource Center.** Related to Star Methods.

**Table S7. Predicted localizations of Chlamydomonas proteins (genome v5.6) by PB-Chlamy.** Related to Figure 7.

(All tables in the attached excel spreadsheet)

**Movie S1. The puncta of Cre15.g640650 show rapid movement.** Related to Figure 2.

## ACKNOWLEDGMENTS

We thank the Princeton University Confocal Microscopy manager Gary Laevsky for instrumentation support; Princeton Institute for Computational Science and Engineering for providing research computing resources; Laurent Cournac, Ben Engel, Ursula Goodenough, Olivier Vallon, Christoph Benning, Ned Wingreen, Xiaobo Li, Silvia Ramundo, Luke C.M. Mackinder, Moritz T. Meyer, Shan He, Alice Lunardon, Jessica H. Hennacy, Sabrina Ergun, Yuliya Zubak, Moshe A. Kafri, Eric Franklin for discussions and feedback on the manuscript; Marie Bao, as part of Life Science Editors, for help with editing the manuscript. The project was funded by DOE grant DE-SC0020195, HHMI/Simons Foundation grant 55108535, and the Lewis-Sigler Scholars Fund. Martin Jonikas is a Howard Hughes Medical Institute Investigator.

---

**Figure S1.**
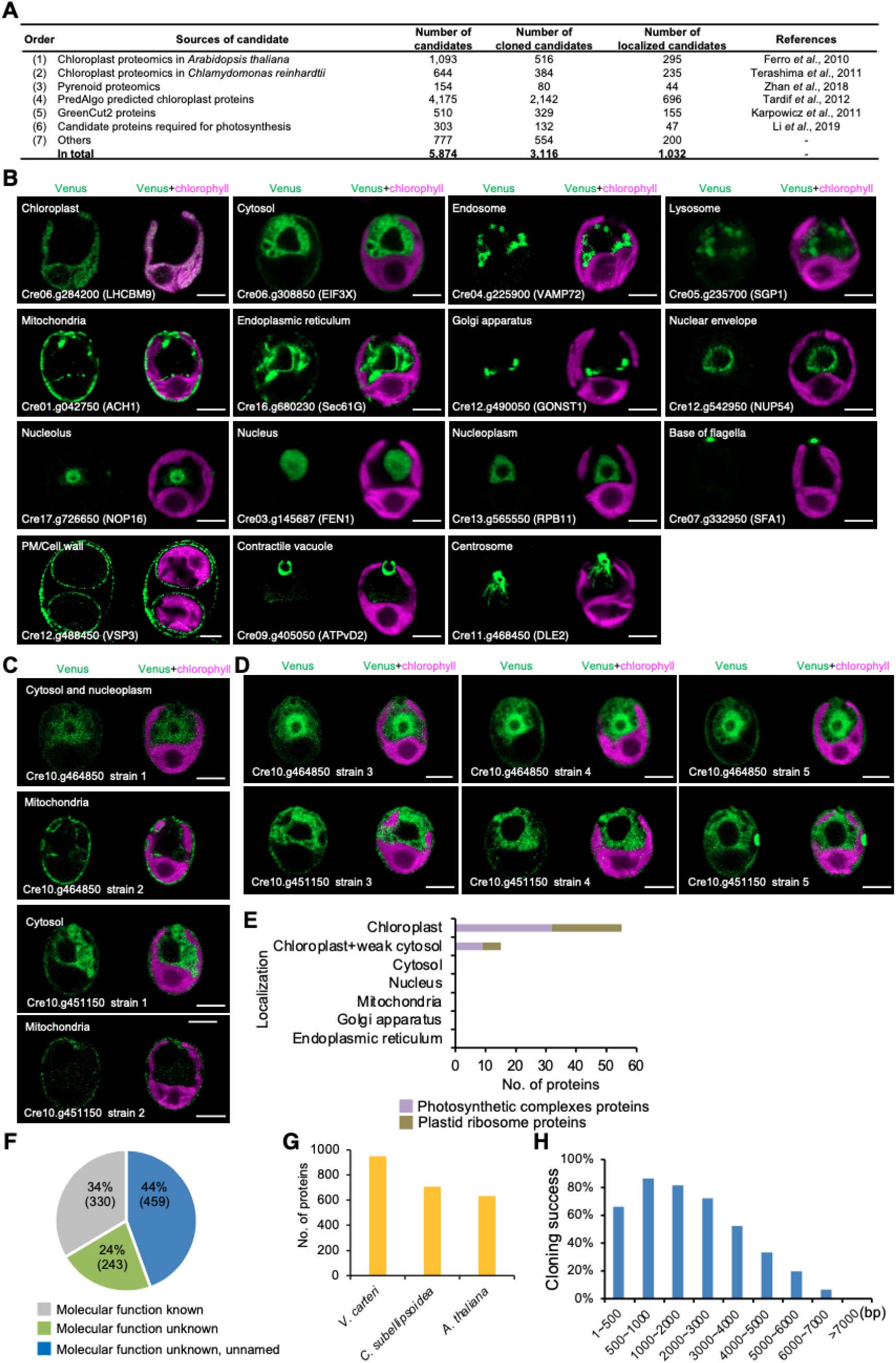
We localized 1,032 proteins from 5,874 target proteins. Related to Figure 1. (A) A summary of seven sources from which target genes were selected. The number of target genes, number of cloned genes, and number of successfully localized proteins are shown. (B) Representative images of organelle marker proteins that localized to different cellular locations. (C) Our localization data showed high reproducibility and chloroplast enrichment of photosynthetic complex proteins and plastid ribosome proteins. Out of 654 proteins for which we obtained two or more independent transformants, the localizations in the independent transformants disagreed for two proteins (<1%), Cre10.g464850 and Cre10.g451150. For each of these two proteins, representative images of the two independent transformants are shown. (D) Representative images of the localization of Cre10.g464850 and Cre10.g451150 in additional transformants, indicating that Cre10.g464850 is localized to cytosol and nucleoplasm; Cre10.g451150 is localized to cytosol. All scale bars are 5 μm. (E) The localization enrichment of plastid ribosome proteins and photosynthetic complexes proteins in chloroplast. (F) Characterization status of our localized proteins. (G) Number of localized proteins conserved in *Volvox carteri*, *Coccomyxa subellipsoidea*, or *Arabidopsis thaliana*. (H) Correlation of cloning success with open reading frame size.

**Figure S2.**
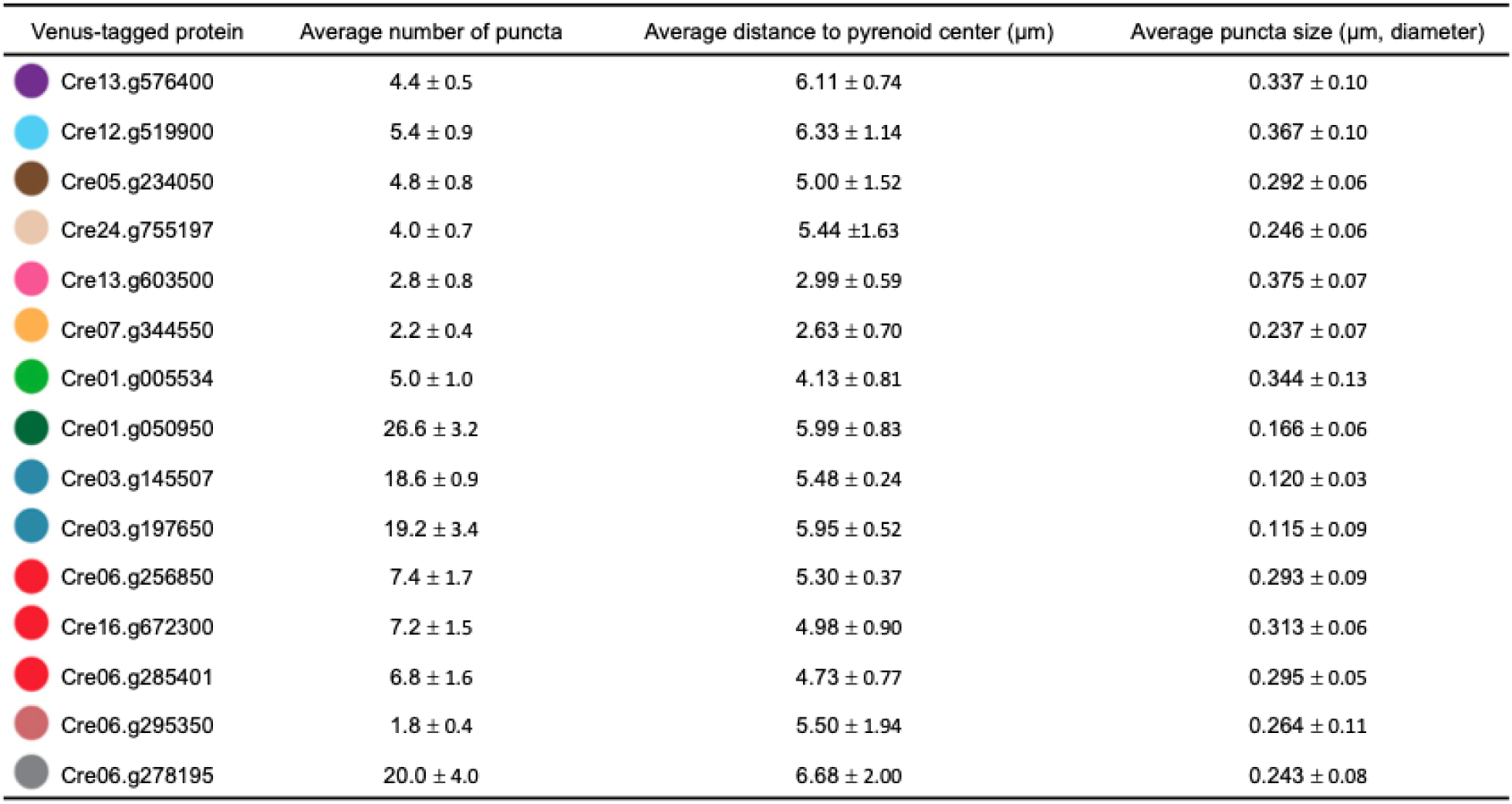
Novel chloroplast punctate structures showed difference in the number of puncta, puncta position, and puncta size. Related to Figure 2. Novel punctate structures showed differences in the average position, number, and size of puncta. For each punctate structure, its mean distance to the pyrenoid center, the average number of puncta, and the mean punctus size are shown. Mean values ± SD from five independent cells are shown. Each structure is assigned a unique color that is used in Figure 2.

**Figure S3.**
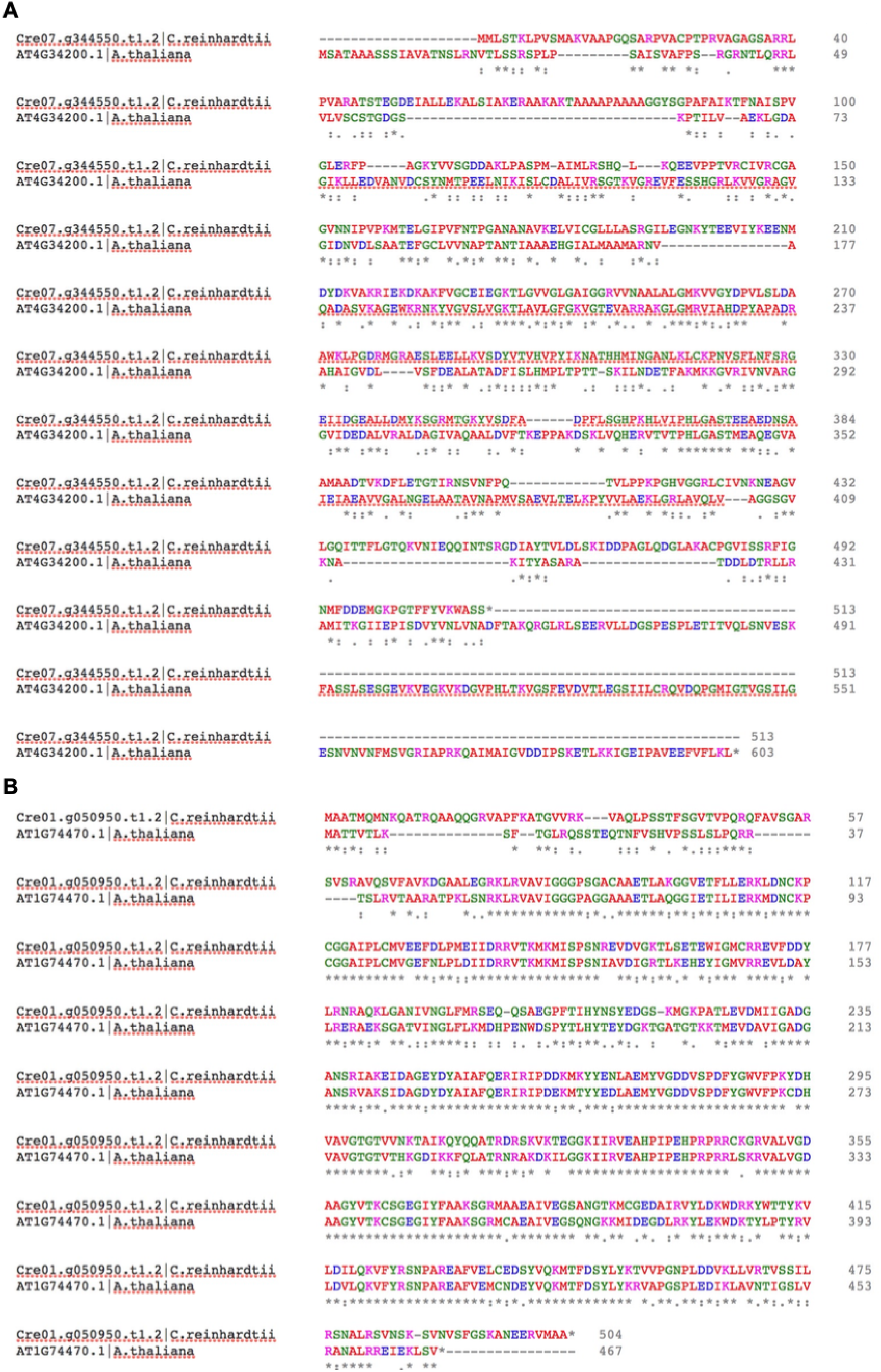
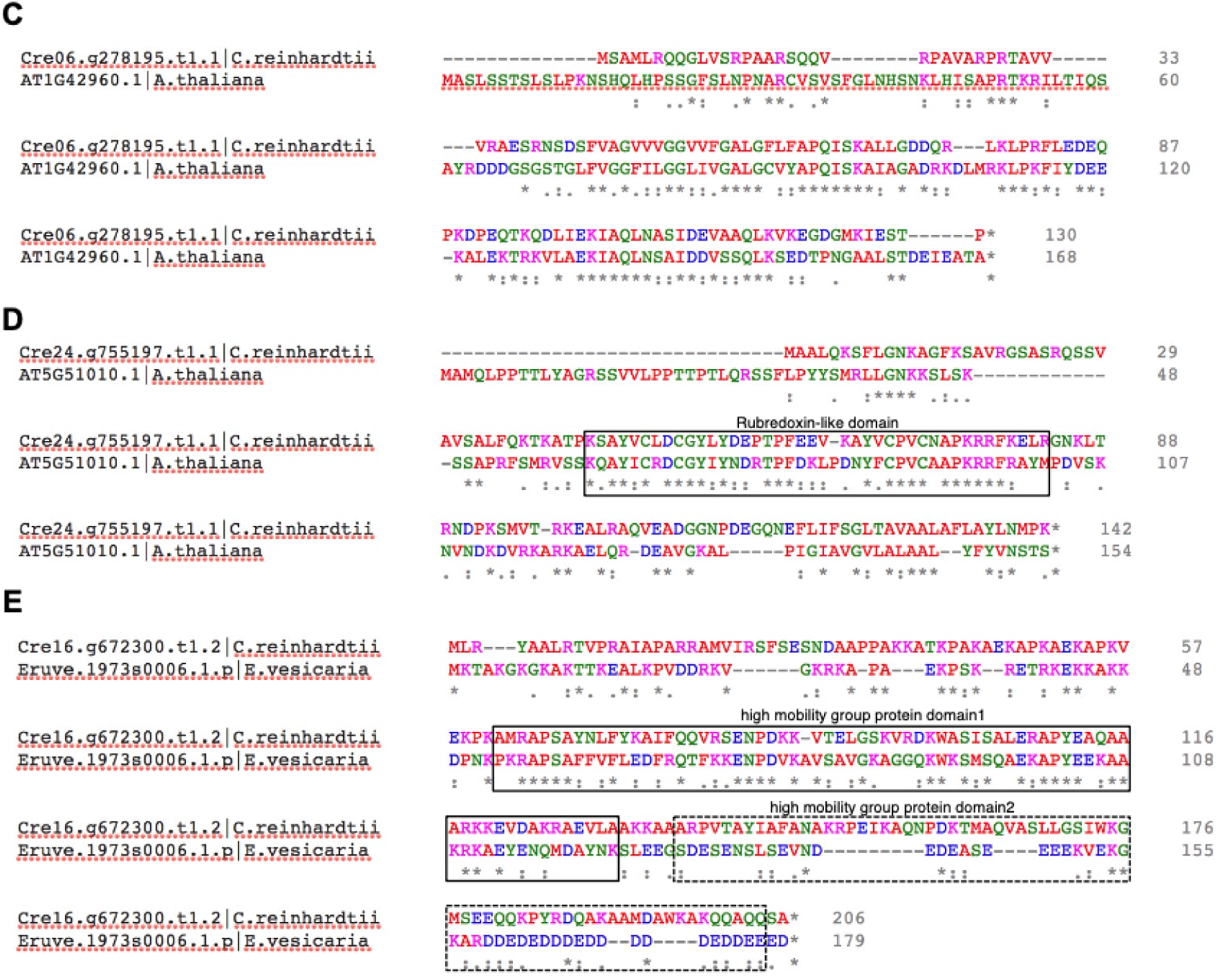
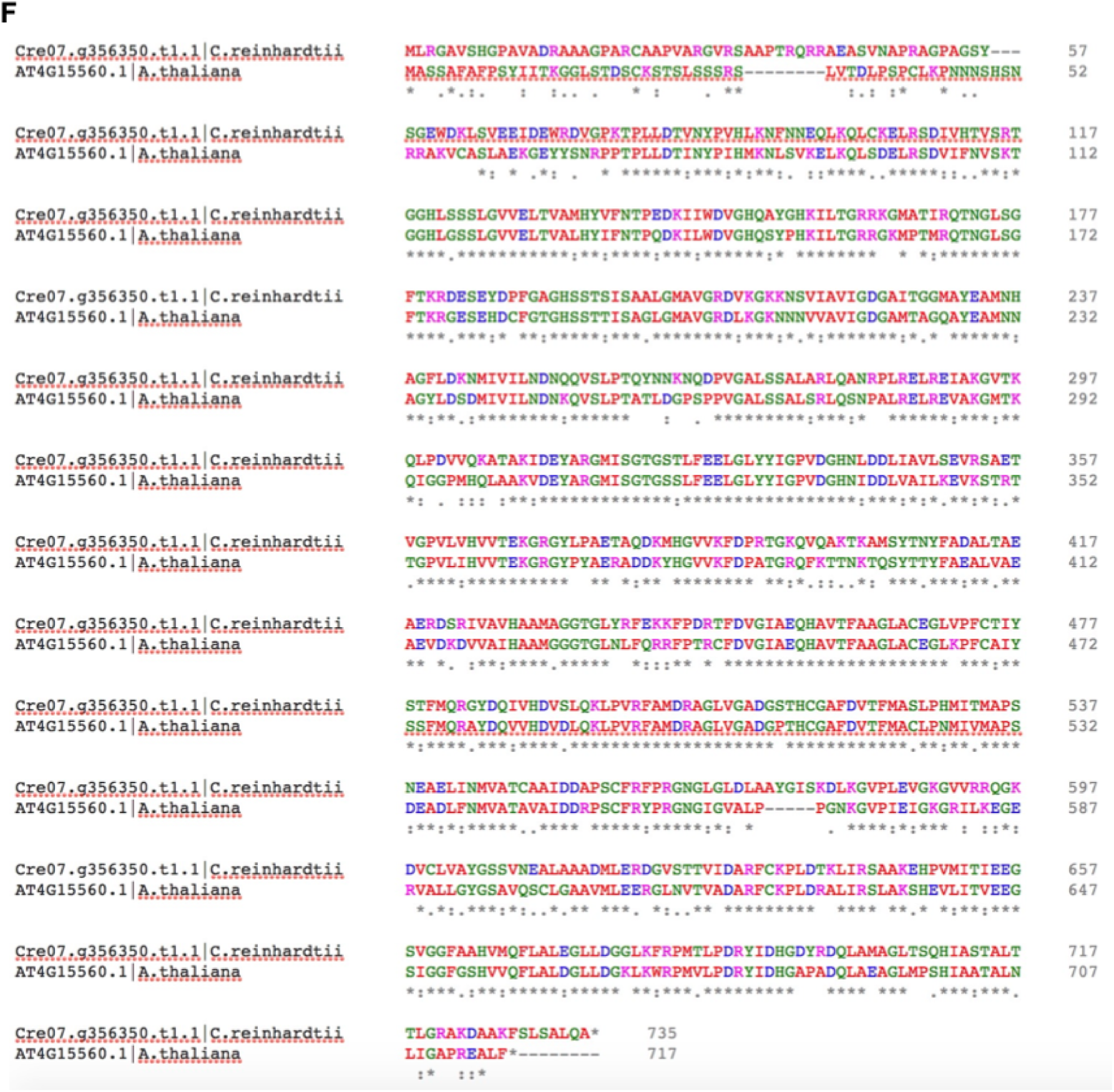
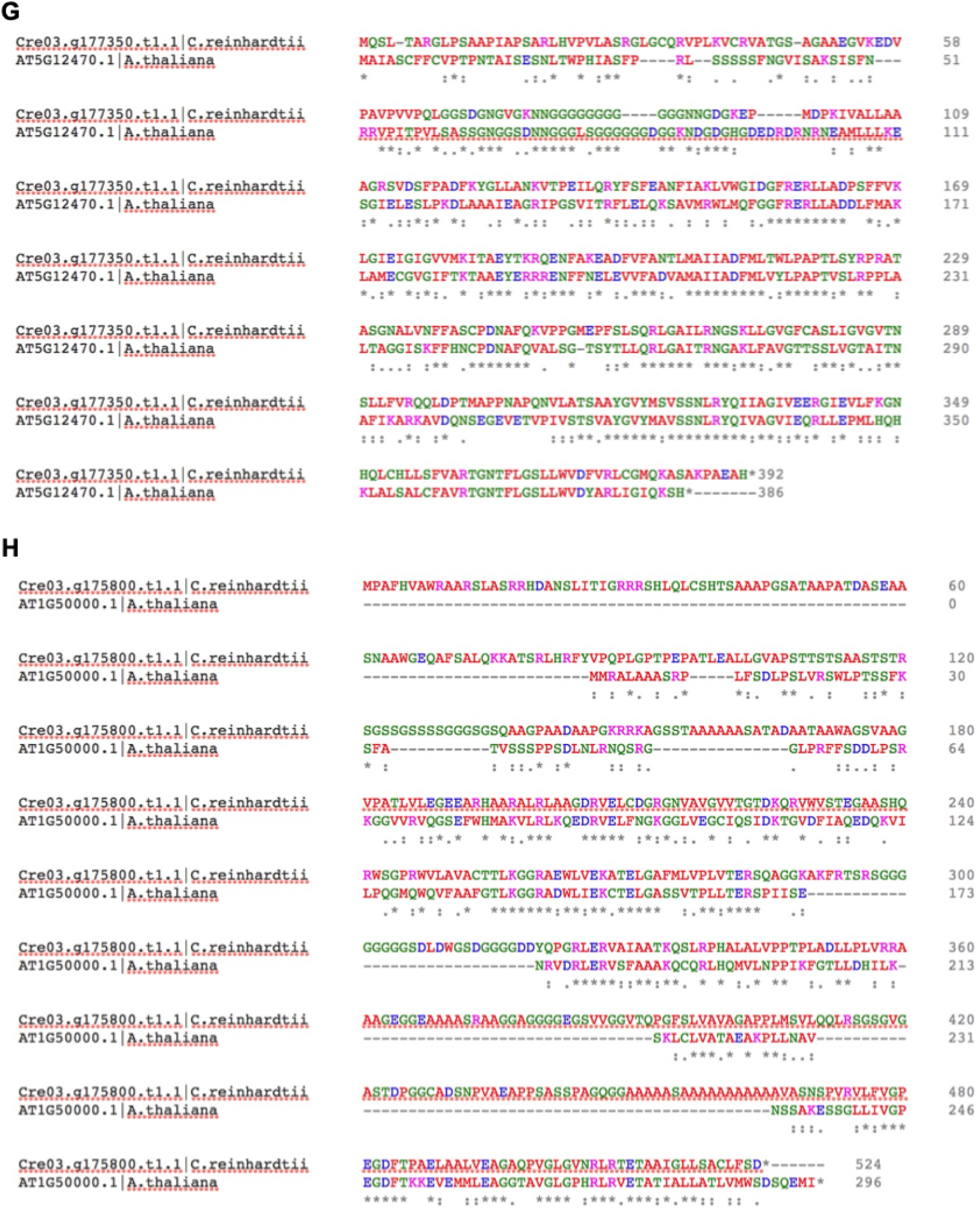
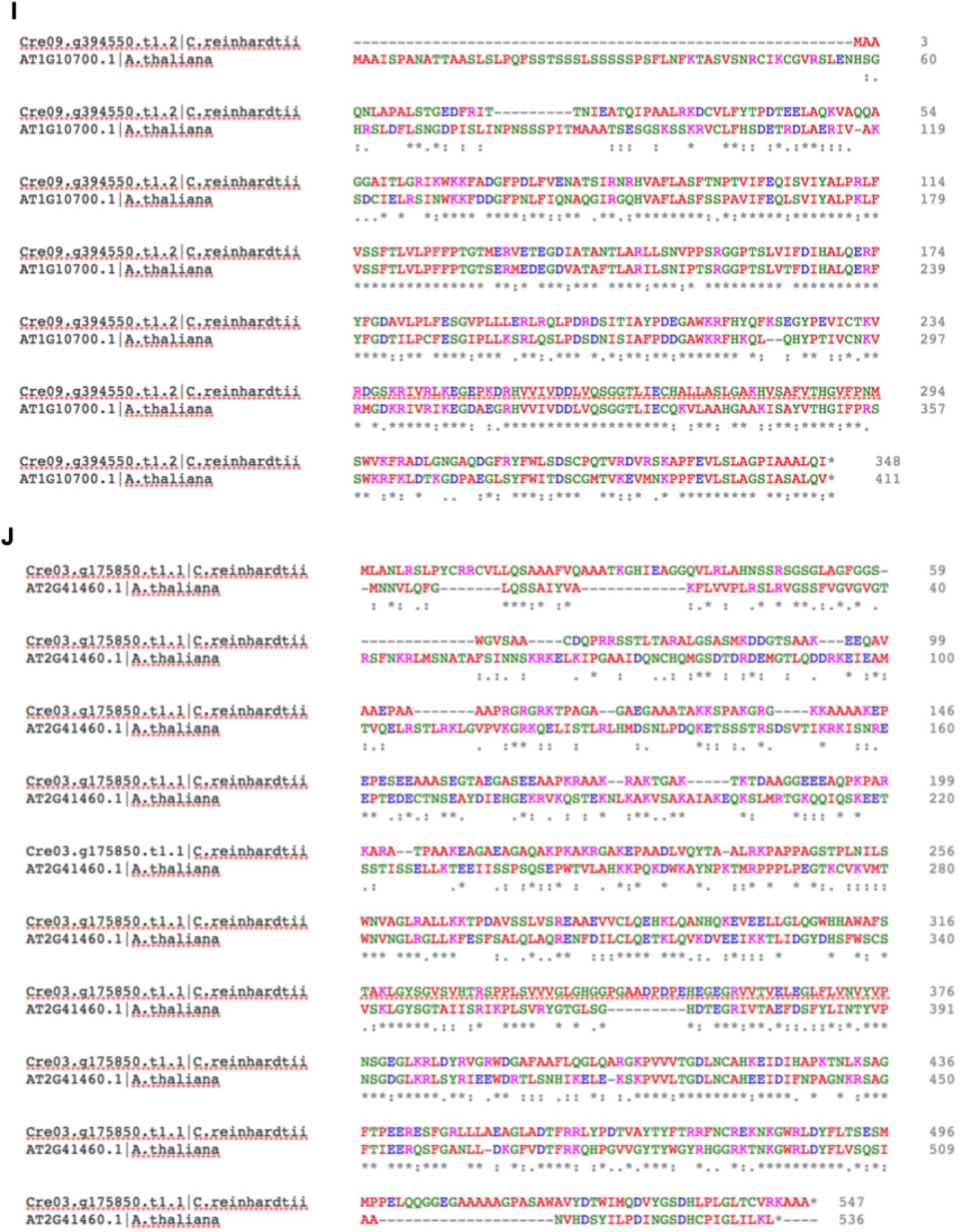
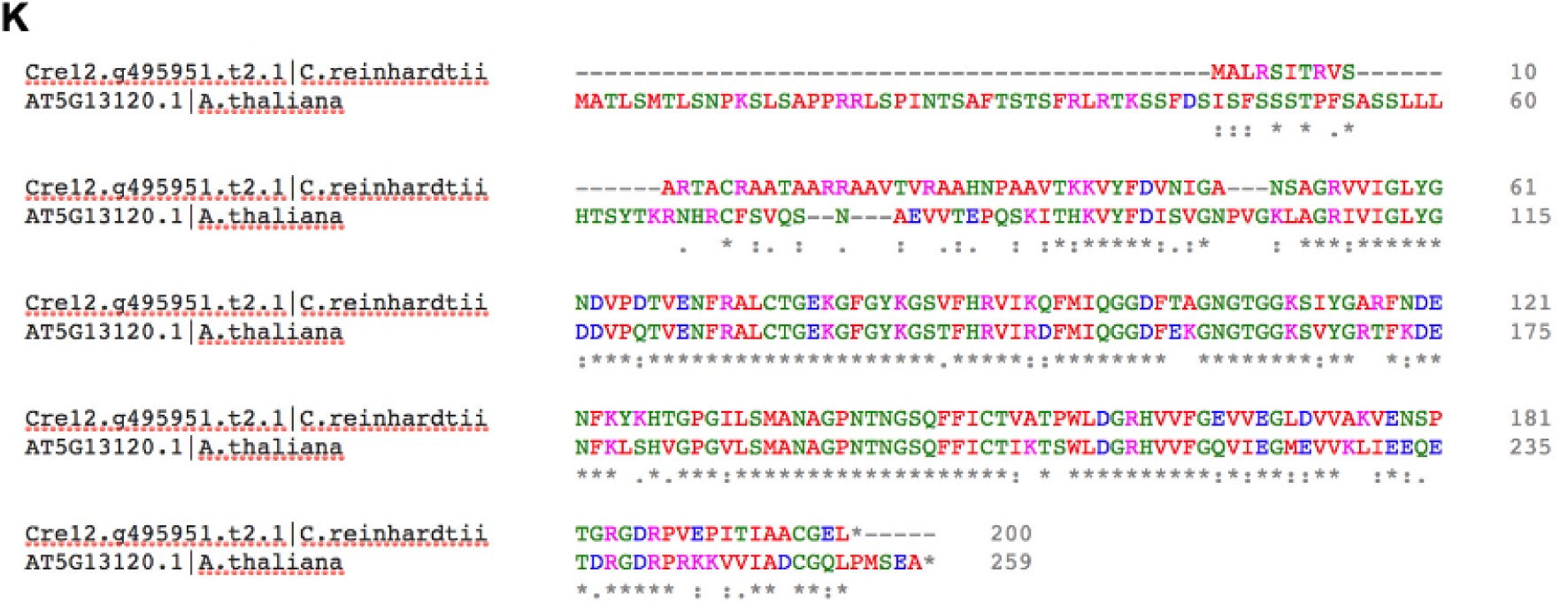
Amino acid alignment of conserved Chlamydomonas proteins with their homologs in land plants. Related to Figure 2-4. The alignment was performed using the Clustal Omega. The colors indicate amino acids with different biochemical properties. (A) Cre07.g344550 and AT4G34200; (B) Cre01.g050950 and AT1G74470; (C) Cre06.g278195 and AT1G42960; (D) Cre24.g755197 and AT5G51010; (E) Cre16.g672300 and Eruve.1973s006; (F) Cre07.g356350 and AT4G15560; (G) Cre03.g177350 and AT5G12470; (H) Cre03.g175800 and AT1G50000; (I) Cre09.g394550 and AT1G10700; (J) Cre03.g175850 and AT2G41460; (K) Cre12.g495951 and AT5G13120.

**Figure S4.**
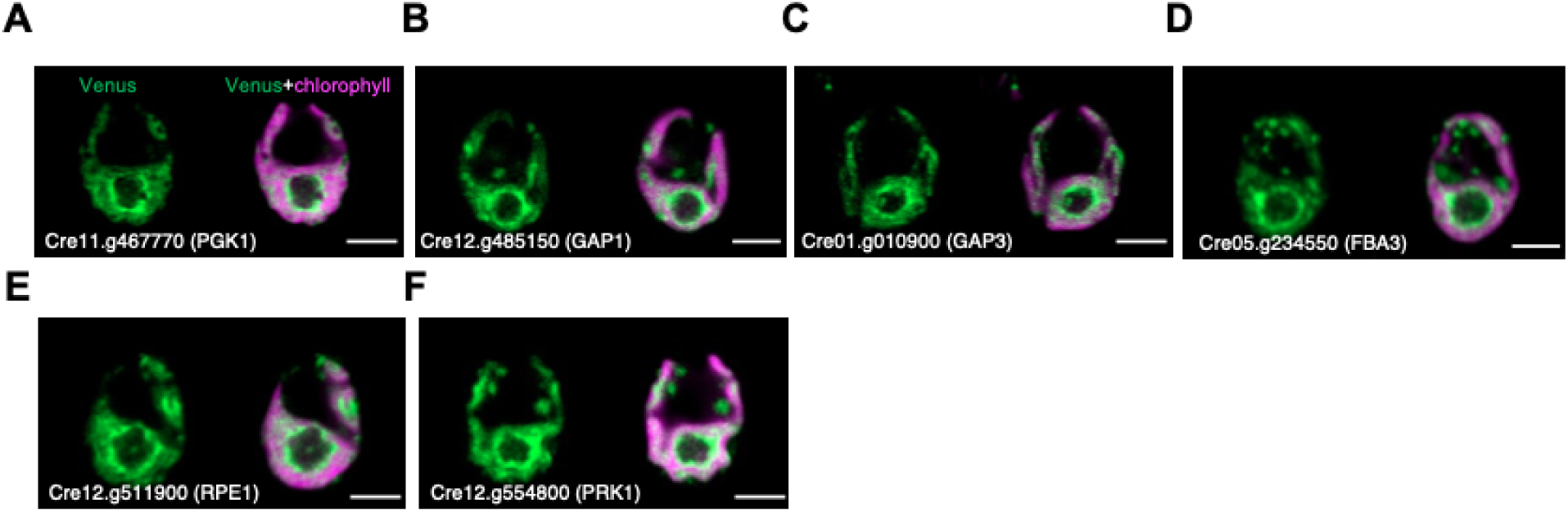
The Calvin-Benson cycle enzymes showed enrichment around pyrenoid. Related to Figure 4. (A-F) Representative images of Cre11.g467770 (PGK1), Cre12.g485150 (GAP1), Cre01.g010900 (GAP3), Cre05.g234550 (FBA3), Cre12.g511900 (RPE1), and Cre12.g554800 (PRK1) showing enrichment in the stroma around pyrenoid. All scale bars are 5 μm.

**Figure S5.**
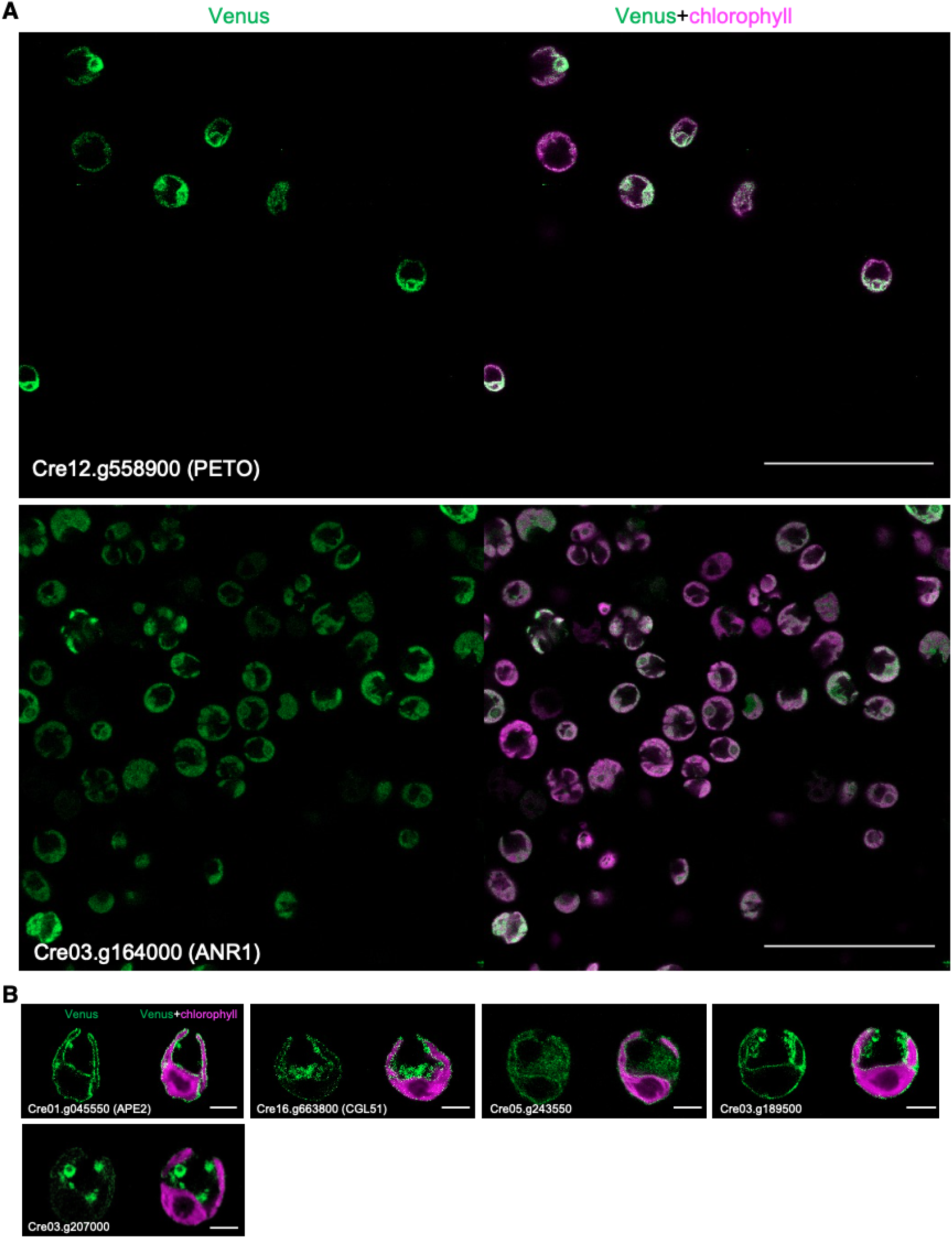
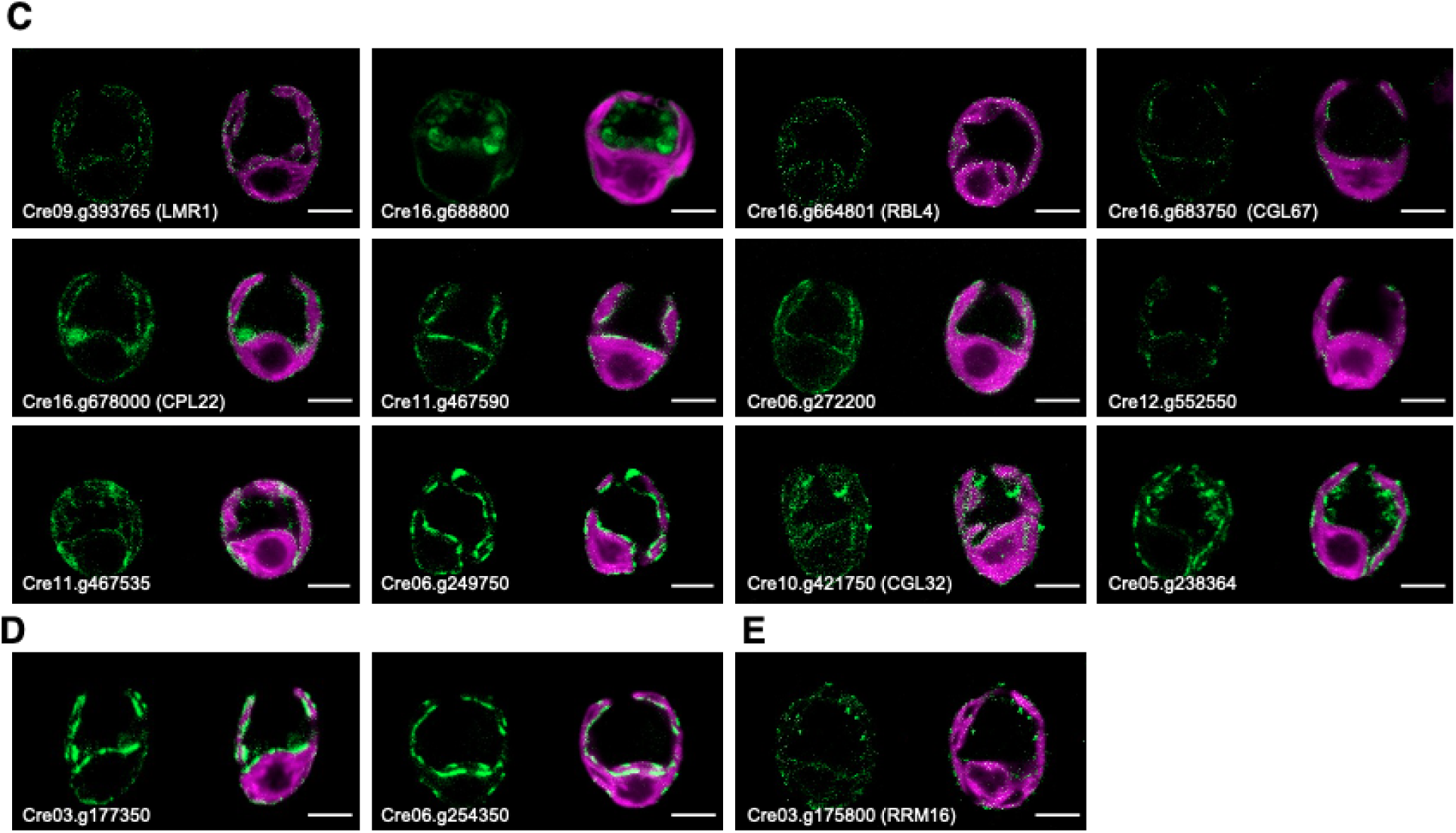
PETO showed different localization patterns from ANR1 and the diverse localizations of proteins localized to chloroplast envelope. Related to Figure 5. (A) Representative localization images of PETO and ANR1. Scale bars, 50 μm. (B) Representative images of proteins homogeneously localized throughout the chloroplast envelope. Scale bars, 5 μm. (C) Representative images of proteins localized to patches along the chloroplast envelope. Scale bars, 5 μm. (D) Representative images of proteins localized to nucleus-facing portion of the chloroplast envelope. Scale bars, 5 μm. (E) Representative images of protein localized as punctate dots along the chloroplast envelope. Scale bars, 5 μm.

**Figure S6.**
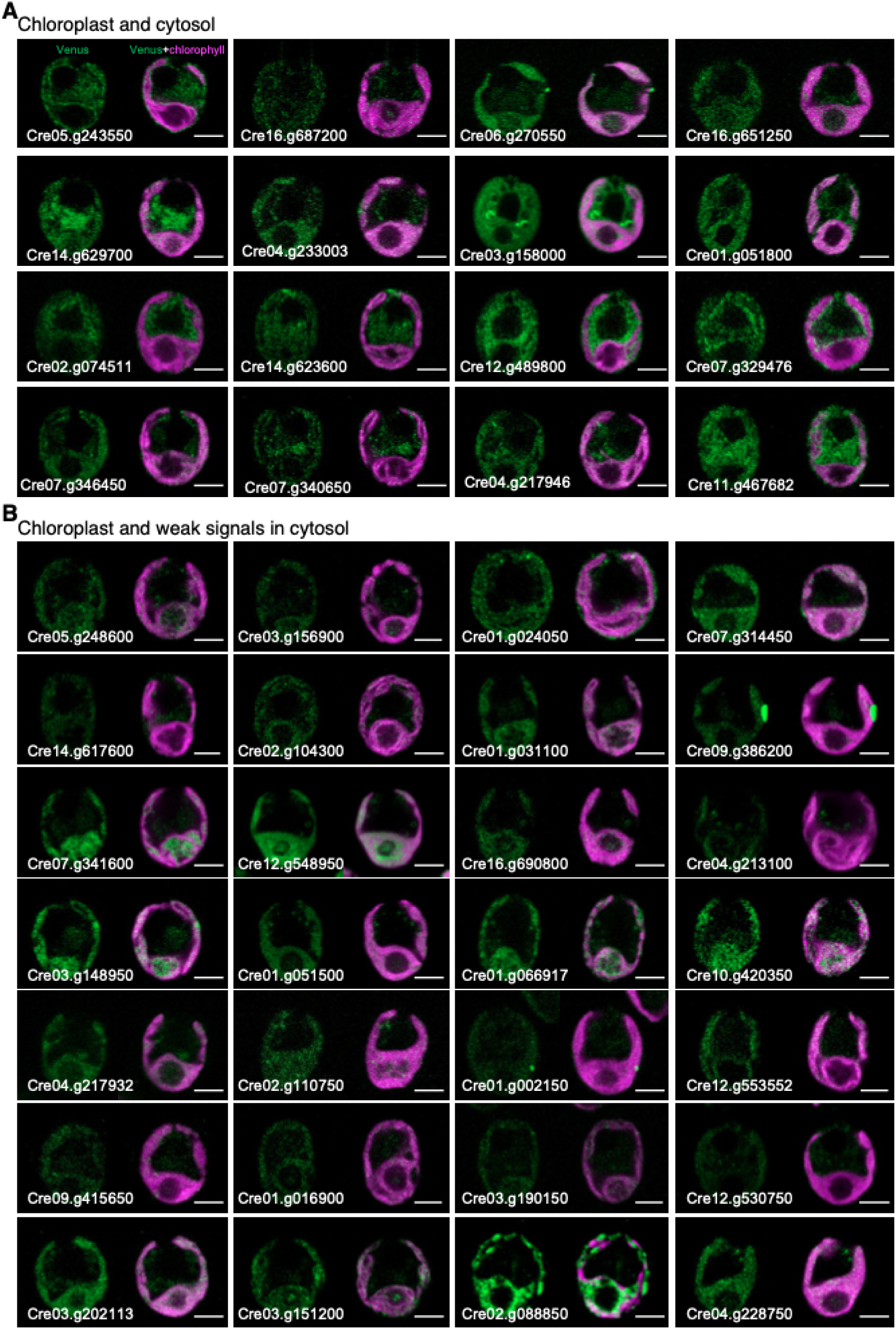
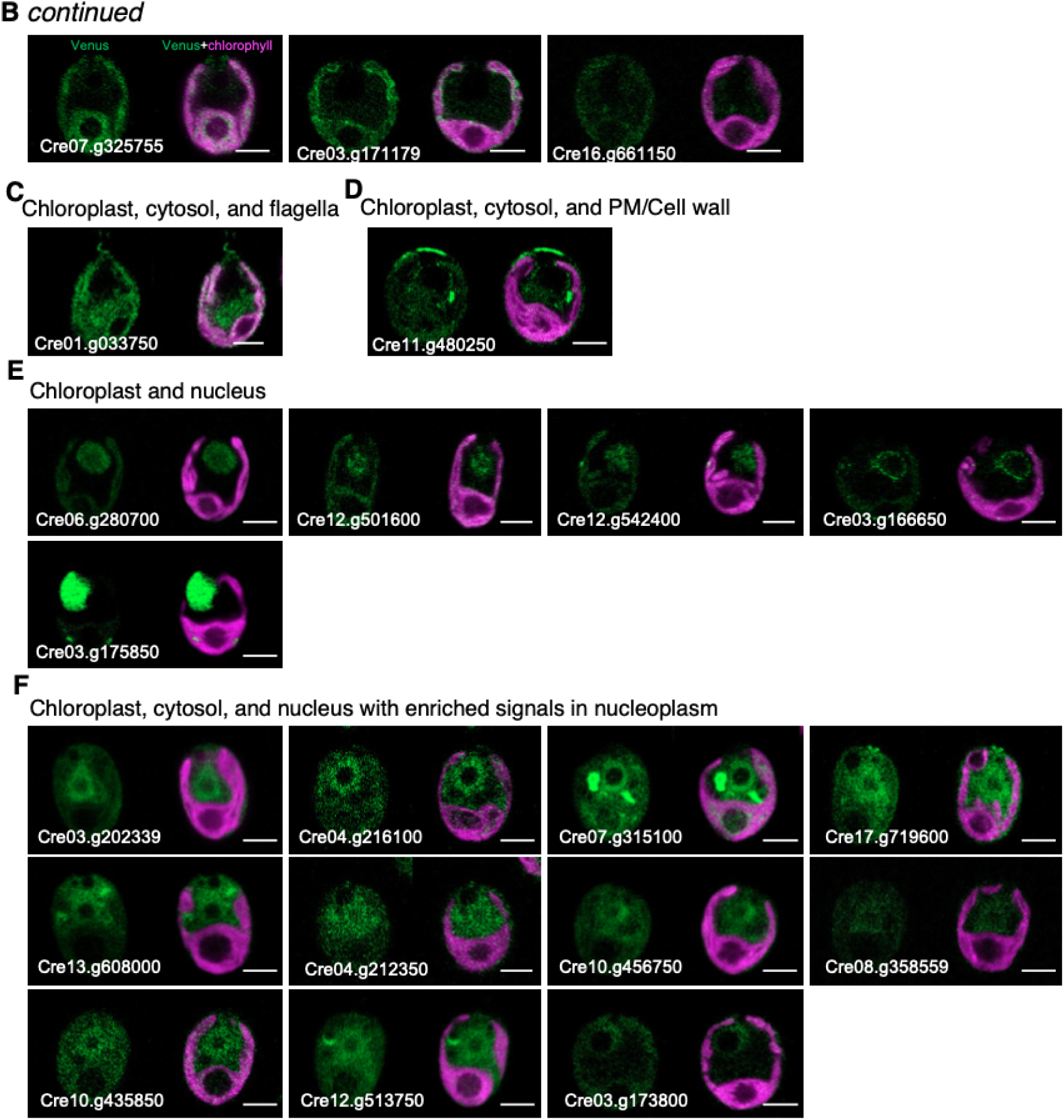
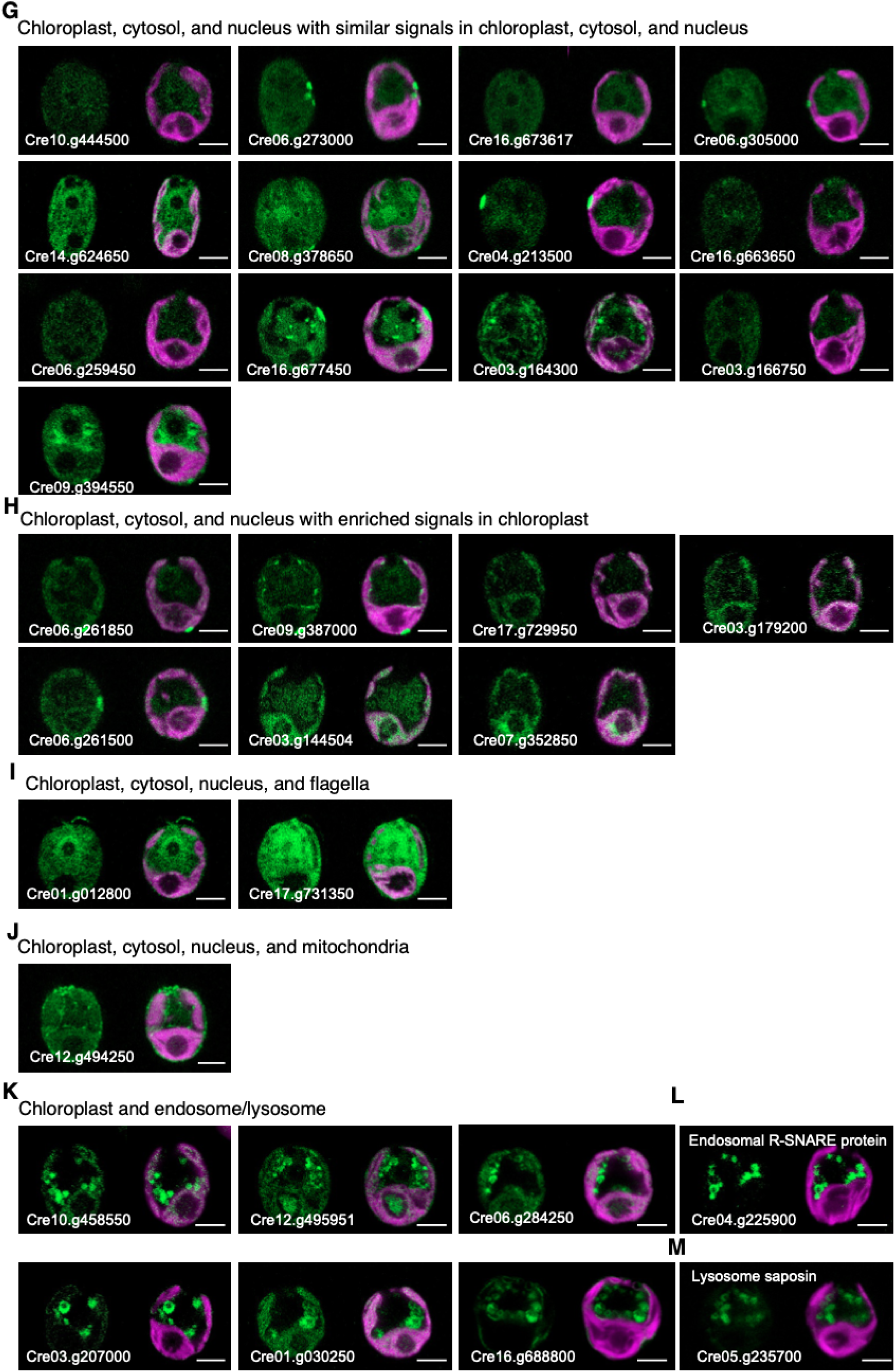
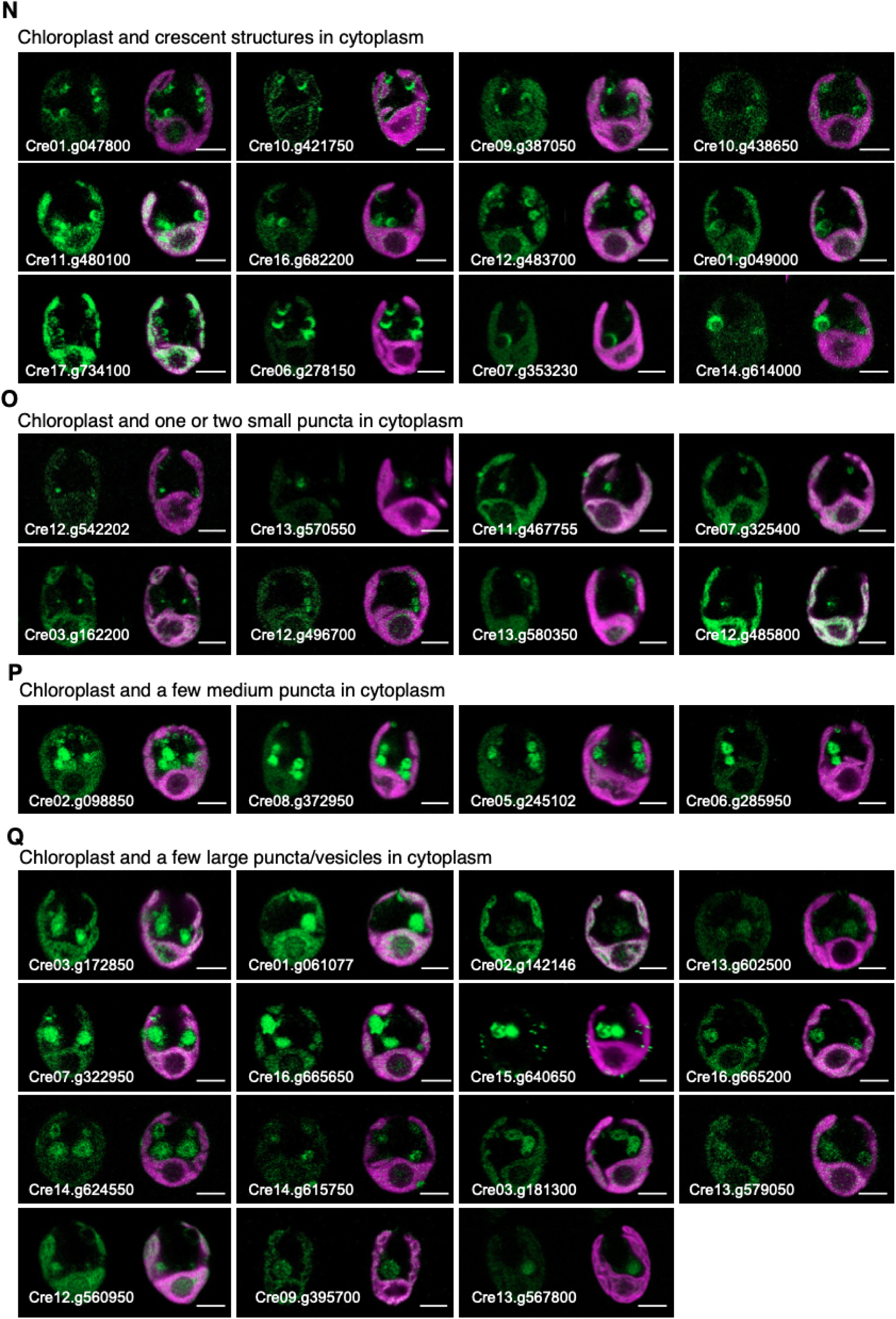
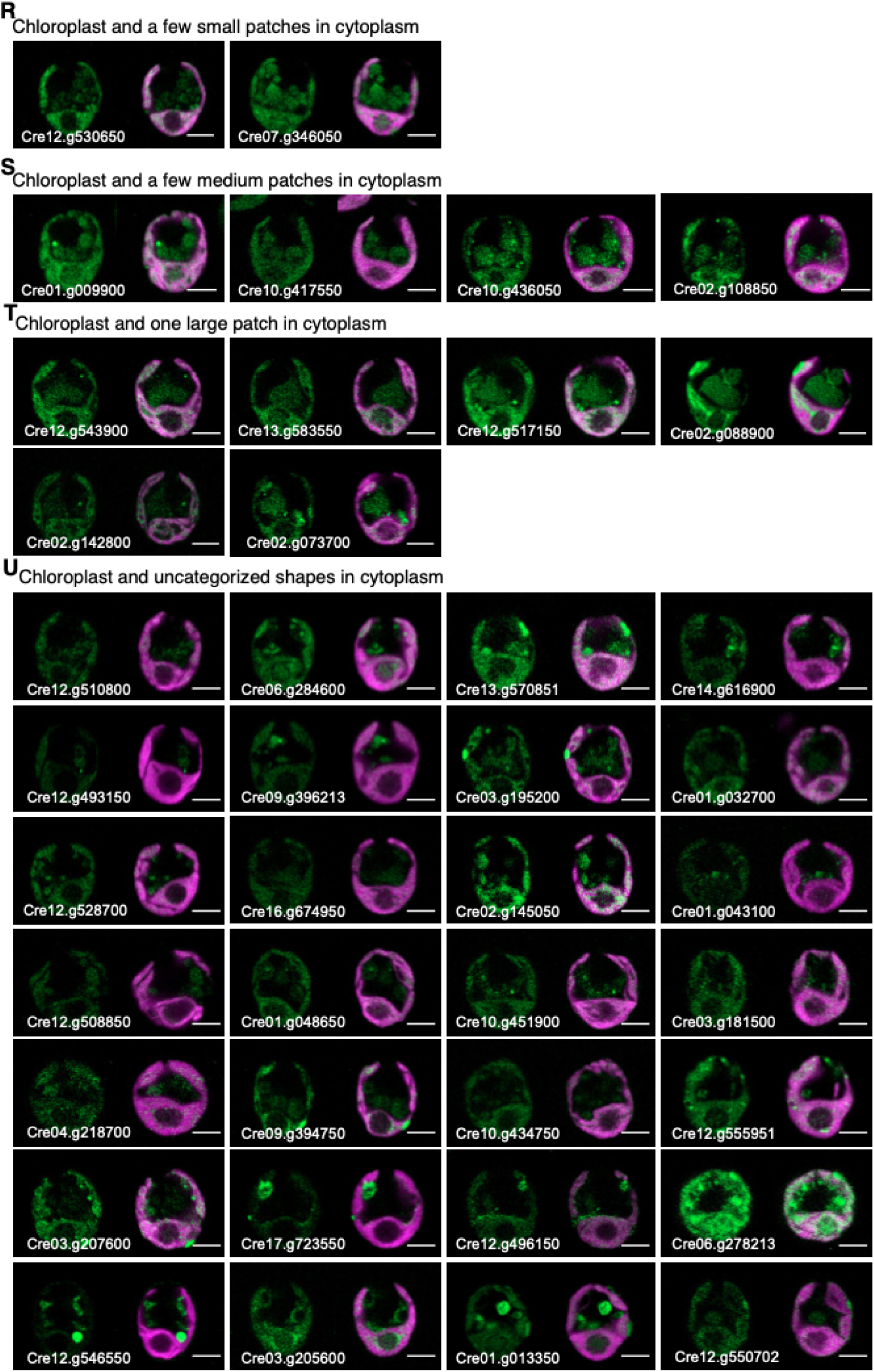
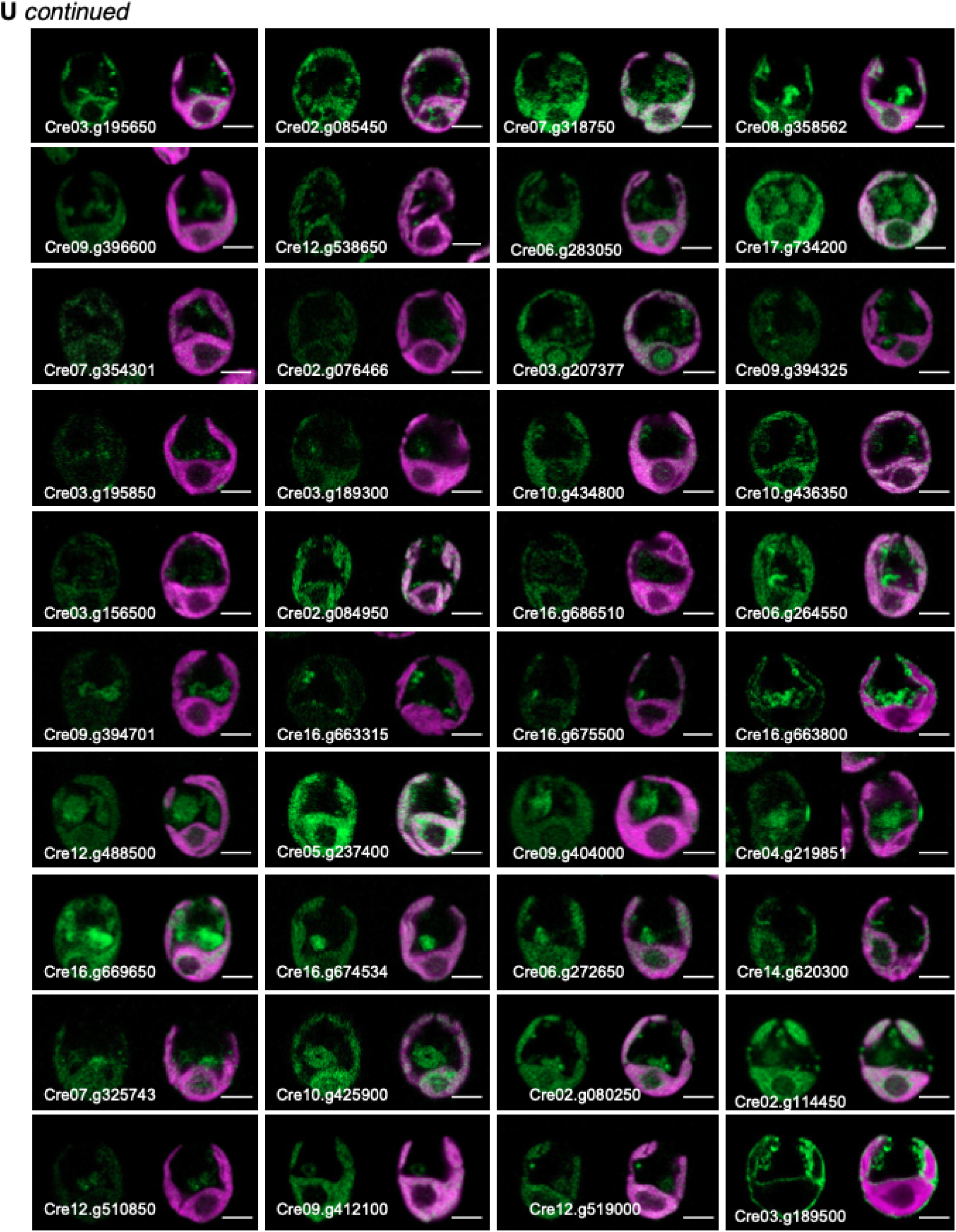
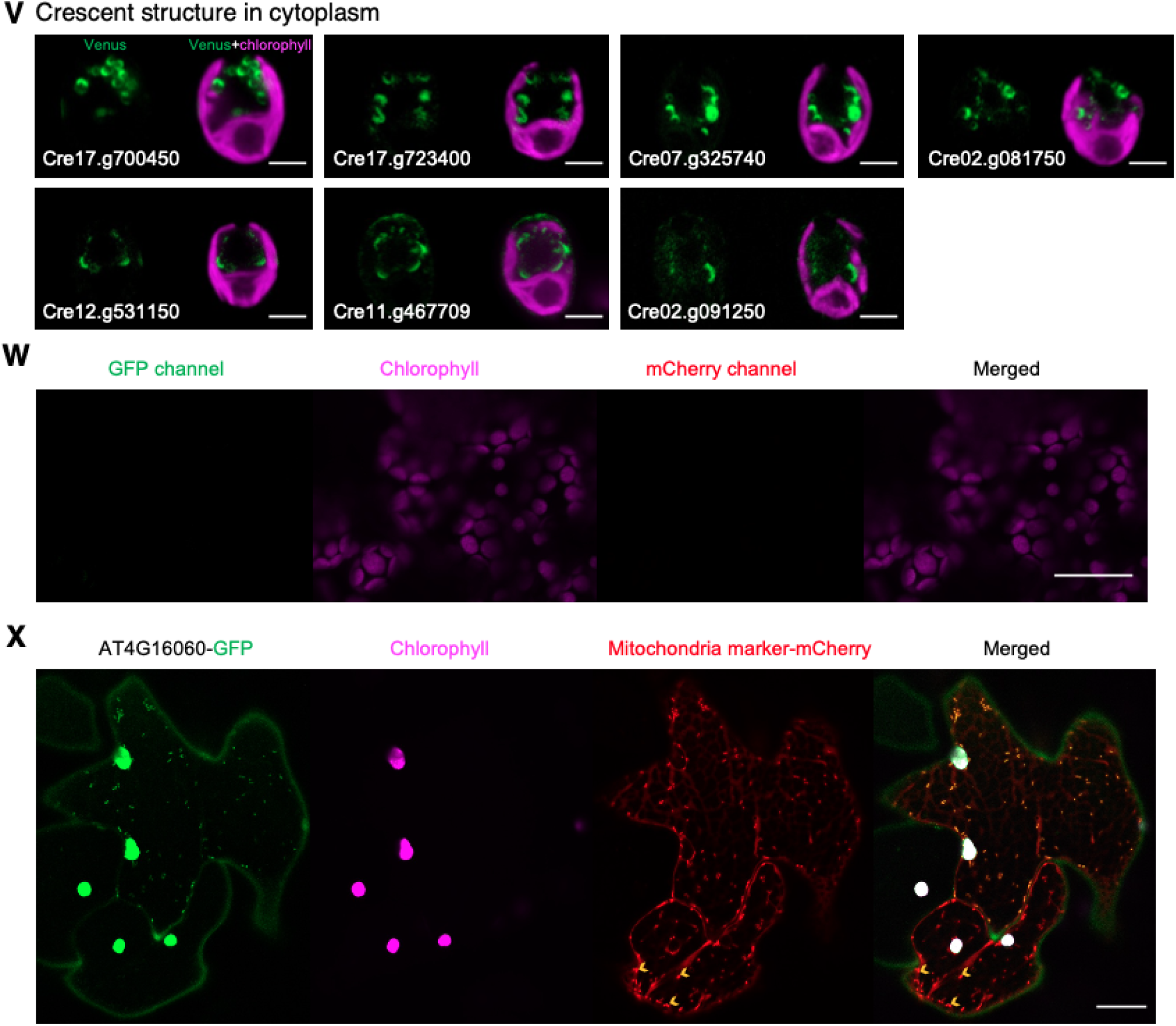
Diverse localizations of proteins localized to chloroplast and other organelles. Related to Figure 6. (A) Representative images of proteins dual-localized to chloroplast and cytosol. (B) Representative images of proteins dual-localized to chloroplast and weak signals in cytosol. (C) Representative images of protein localized to chloroplast, cytosol, and flagella. (D) Representative images of protein localized to chloroplast, cytosol, and PM/Cell wall. (E) Representative images of proteins dual-localized to chloroplast and nucleus. (F) Representative images of proteins localized to chloroplast, nucleus, and cytosol with enriched fluorescence signals in nucleoplasm. (G) Representative images of proteins localized to chloroplast, nucleus, and cytosol with similar fluorescence signals between chloroplast and nucleus. (H) Representative images of proteins localized to chloroplast, nucleus, and cytosol with enriched fluorescence signals in chloroplast. (I) Representative images of proteins localized to chloroplast, nucleus, cytosol, and flagella. (J) Representative images of proteins localized to chloroplast, nucleus, cytosol, and mitochondria. (K) Representative images of proteins dual-localized to chloroplast and endosomes/lysosomes (many small puncta or vesicles in cytoplasm). (L) Representative images of endosomal R-SNARE protein localized to endosomes. (M) Representative images of lysosome saposin localized to lysosomes. (N) Representative images of proteins dual-localized to chloroplast and crescent structures in the cytoplasm. (O) Representative images of proteins dual-localized to chloroplast and one or two small puncta in the cytoplasm. (P) Representative images of proteins dual-localized to chloroplast and a few medium puncta in the cytoplasm. (Q) Representative images of proteins dual-localized to chloroplast and a few large puncta/vesicles in the cytoplasm. (R) Representative images of proteins dual-localized to chloroplast and a few small patches in the cytoplasm. (S) Representative images of proteins dual-localized to chloroplast and a few medium patches in the cytoplasm. (T) Representative images of proteins dual-localized to chloroplast and one large patch in the cytoplasm. (U) Representative images of proteins dual-localized to chloroplast and uncategorized shapes in the cytoplasm. (V) Representative images of proteins exclusively to the crescent structures in the cytoplasm. All scale bars in (A)-(V) are 5 μm. (W) Representative images of chloroplast in the tobacco leaf cells without infiltration with Agrobacterium tumefaciens. (X) Representative images of mitochondria mCherry marker (CD3-991) in the tobacco leaf cell without GFP expression. The yellow arrows indicate mitochondria. All scale bars in (W) and (X) are 20 μm.

**Figure S7.**
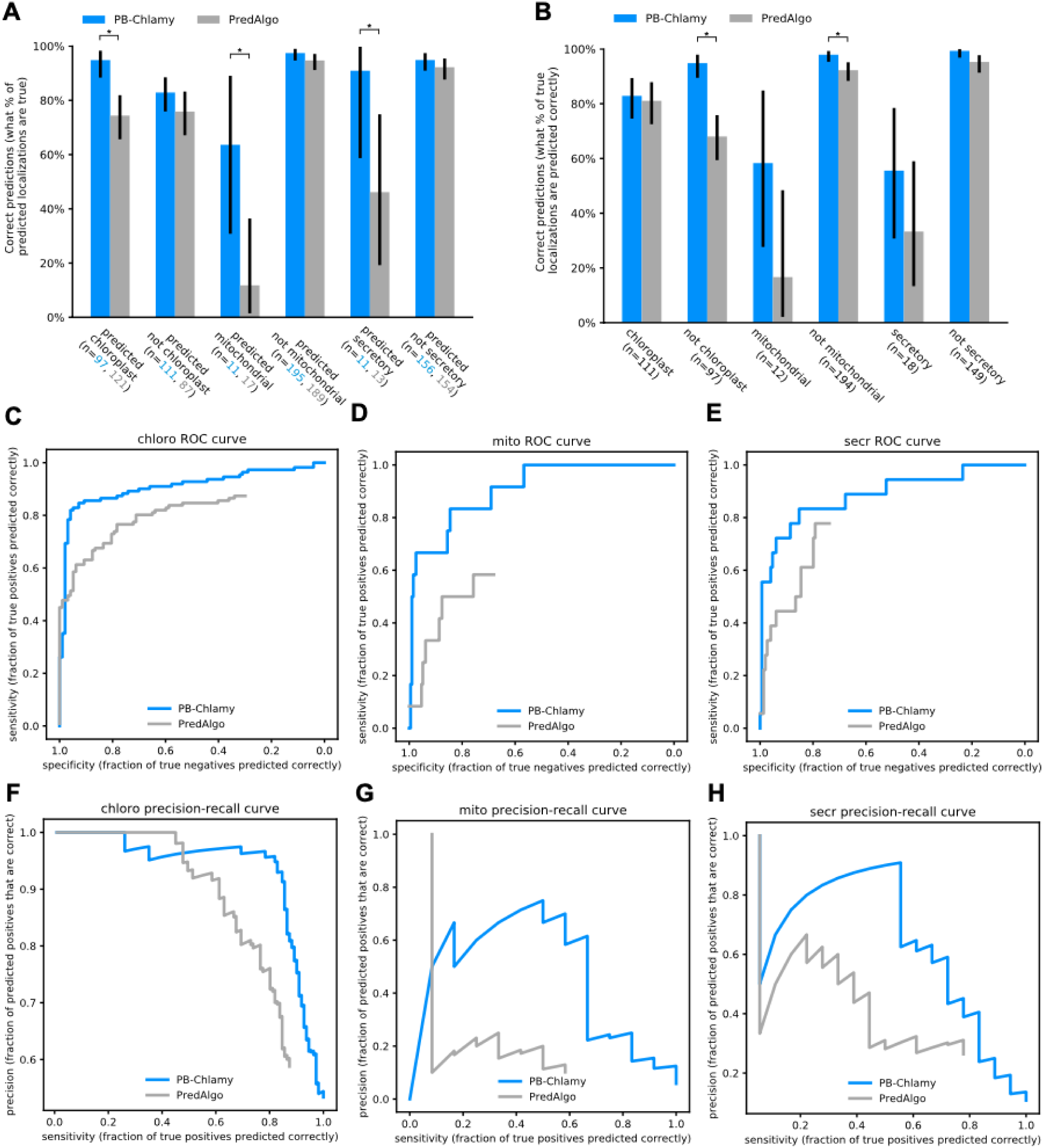
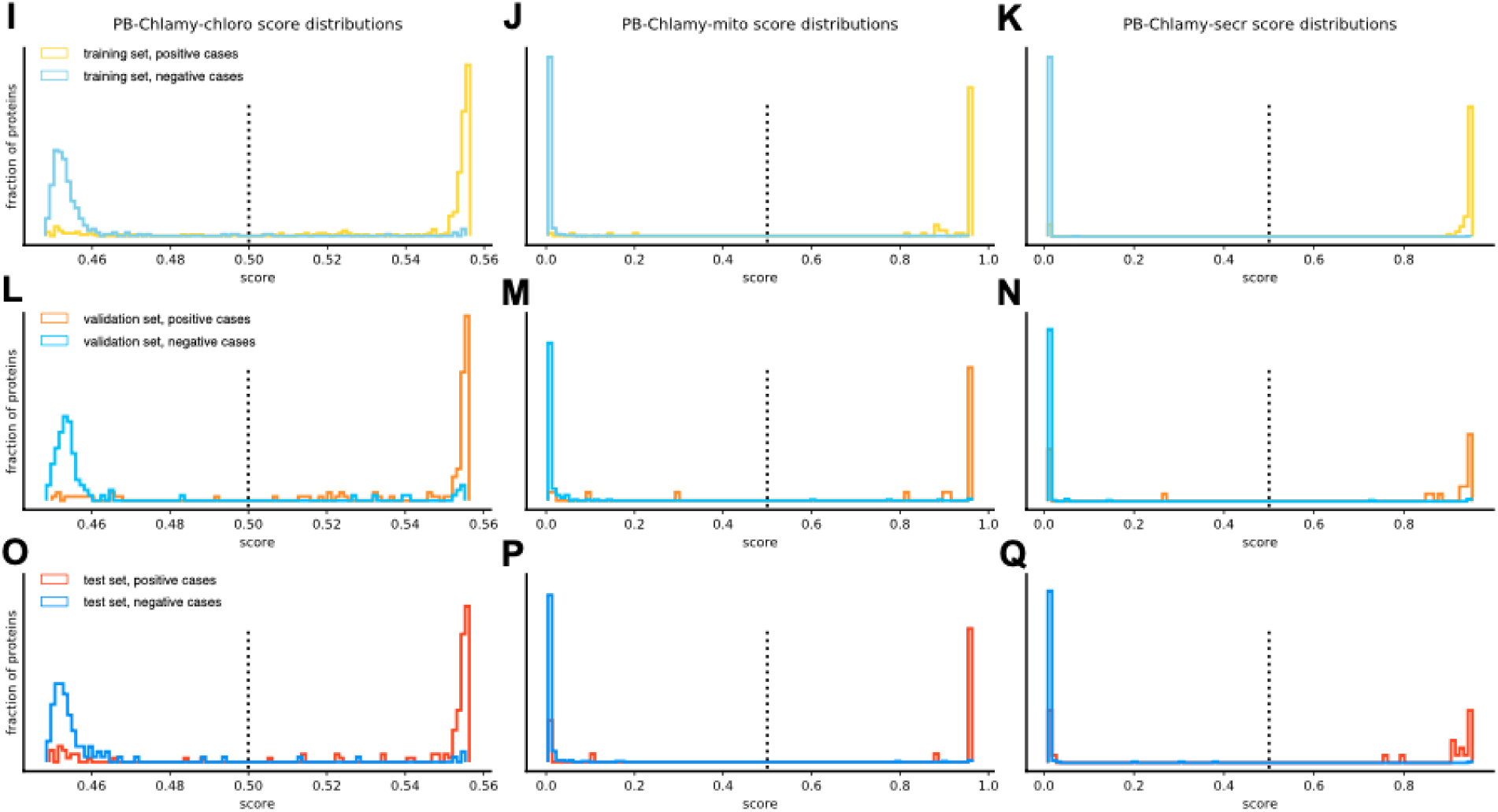
PB-Chlamy reliably predicts Chlamydomonas protein localizations. Related to Figure 7. (A) The percent of predicted positive and negative protein localizations that are correct, for PB-Chlamy and PredAlgo. Error bars represent binomial 90% confidence intervals. The number of proteins varies between PB-Chlamy and PredAlgo, as the two programs predict different numbers of positive and negative proteins. The p-values comparing PB-Chlamy and PredAlgo (using Fisher’s exact test) are: chloroplast 0.000036, non-chloroplast 0.28, mitochondrial 0.010, non-mitochondrial 0.20, secretory 0.033, non-secretory 0.36. The asterisk * indicates p<0.05. (B) The percent of true positive and negative protein localizations that were correctly classified by PB-Chlamy and PredAlgo. Error bars represent binomial 90% confidence intervals. The p-values comparing PB-Chlamy and PredAlgo using Fisher’s exact test are: chloroplast 0.86, non-chloroplast 0.0000016, mitochondrial 0.089, non-mitochondrial 0.016, secretory 0.31, non-secretory 0.067. The asterisk * indicates p<0.05. (C-E) ROC curves comparing PB-Chlamy and PredAlgo for each localization category. PredAlgo uses a static score cutoff for predictions, therefore ROC curves terminate at this cutoff and do not reach a sensitivity of 1. (F-H) Precision-recall curves comparing PB-Chlamy and PredAlgo for each localization category. (I-Q) The raw PB-Chlamy prediction score distributions, normalized, for each chloroplast (I, L, O), mitochondrial (J, M, P) and secretory (K, N, Q), showing the training (I, J, K), validation (L, M, N) and test (O, P, Q) sets, with true positives and true negatives overlaid. The dotted bar at 0.5 is the cutoff for positive vs negative predictions.

## Notes

https://www.chlamylibrary.org/allTaggedStrains

### Competing Interest Statement

The authors have declared no competing interest.

